# Oligodendroglial precursors orchestrate immune network instigating early demyelination in experimental autoimmune encephalomyelitis

**DOI:** 10.1101/2023.10.10.561804

**Authors:** Qi Wang, Taida Huang, Zihan Zheng, Yixun Su, Zhonghao Wu, Guangdan Yu, Yang Liu, Xiaorui Wang, Hui Li, Xiaoying Chen, Zhuoxu Jiang, Jinyu Zhang, Yuan Zhuang, Yi Tian, Qingwu Yang, Alexei Verkhratsky, Ying Wan, Chenju Yi, Jianqin Niu

## Abstract

The immunomodulatory cellular network that triggers early inflammation and demyelination, the key steps in multiple sclerosis (MS) pathogenesis remains poorly characterized. Here, we demonstrate that overactivation of Wnt pathway promotes pathological transformation of oligodendrocyte precursor cells (OPCs) to replicate pathological OPCs in human MS. In mouse experimental autoimmune encephalomyelitis (EAE), pathological OPCs attract CD4^+^ T-helper 1 (Th1) cells into the spinal cord and brain through CC-chemokine ligand 4 (CCL4), whilst OPCs cooperate with Th1 cells inducing transformation of cytotoxic macrophages that execute early demyelination. Simultaneously, Th1 cells and cytotoxic macrophages upregulate Wnt signaling and CCL4 expression in OPCs, thus exerting positive feedback onto the OPC-immune cascade and establishing a vicious cycle propagating EAE pathogenesis. Breaking this cascade by targeting CCL4 reduces immune cell infiltration, alleviates demyelination, and attenuates EAE severity. Our findings demonstrate a closely coordinated network of OPCs and immune cells therefore providing an alternative insight into MS pathophysiology.

## Introduction

To date, it is universally accepted that multiple sclerosis (MS) is instigated by autoreactive immune cells that reside within and infiltrate the central nervous system (CNS) ^1,2^, while several types of immune cells are implicated in causing progressive demyelination and axonal damage ^1,3^. However, which cell type(s) define(s) early pathogenesis remains debatable ^4-7^, thus impeding development of early intervention strategies for MS containment before irreversible neuropathological outcomes.

Early upregulation of Wnt signaling is observed in several inflammation-associated diseases ^8^, including MS ^9^. In these diseases, Wnt signaling may perform an essential role in the regulation of immune cells ^10^. Recently we identified an overactivated-Wnt pathway in oligodendrocyte precursor cells (OPCs) from human MS brain samples ^11,12^, dovetailing with the emerging evidence of OPCs acting as ‘immunomodulatory agents’ in MS ^13-15^, and OPC numbers increasing in peri-plaque white matter ^16,17^, at active lesion sites and in undamaged white matter ^18^, where signs of inflammation are detected ^19,20^. Overactivated Wnt arguably induces new, pathological OPCs, which we define as high Wnt activation-OPCs (hWnt-OPCs). Nevertheless, it is unclear whether and how hWnt-OPCs are involved in an immunomodulatory network and participate in demyelination at the onset of MS.

Available MS animal models do not fully recapitulate pathophysiology of the human MS. Our previous studies revealed that OPCs undergo more severe pathological changes in MS patients than that in mouse models, which coincides with much higher levels of Wnt signaling in human postmortem tissues ^11,12^. In the present study, we employed an OPC-specific inducible Wnt-activation mouse strain ^11,12^, which allows in-depth exploration of the immune-modulatory network in a mouse experimental autoimmune encephalomyelitis (EAE) model at a higher Wnt background tone favoring formation of pathological OPCs ^21^. Using this model, we discovered a previously unrecognized pathological hWnt-OPCs-immune cellular network, which provides an alternative insight into the pathogenesis and pathophysiology of MS, and may open a new avenue for development of early protective interventions.

## Results

### hWnt-OPCs accelerate and exacerbate early EAE and promote CD4^+^ T cell recruitment

To determine whether hWnt-OPCs contribute to the early EAE pathogenesis, we firstly examined the correlation between the OPCs pathology and EAE early progression (early-onset stage: 9-day to 14-day post-immunization (dpi); post-onset stage: 15 to 18 dpi, which is reflected by the clinical scores) **(Figure 1A)**. As shown by the sternberger monoclonal-incorporated antibody 32 (SMI32)-labeled non-phosphorylated neurofilaments and myelin basic proteins (MBP)-labeled myelin structures, early signs of EAE, including demyelinating and axonal injury, occurred at 14 dpi and became further aggravated at 18 dpi **(Figure 1B, S1A)**. The number of PDGFRα^+^ OPCs increased in parallel with accumulation of CD4^+^ T, but not CD8^+^ T cells or CD19^+^ B immune cells **(Figure 1C, D)**. At the same time, ring finger protein 43 (RNF43), a newly identified indicator of Wnt activation in hWnt-OPCs in MS ^12^, was upregulated in the early stage of EAE progression **(Figure 1E)**. At a later stage of EAE, at 28 dpi, the number of OPCs started to decline together with decrease in CD4^+^ T cells, parallel with other immune cells also descending in the lesioned areas **(Figure S1B)**.

**Figure 1.**
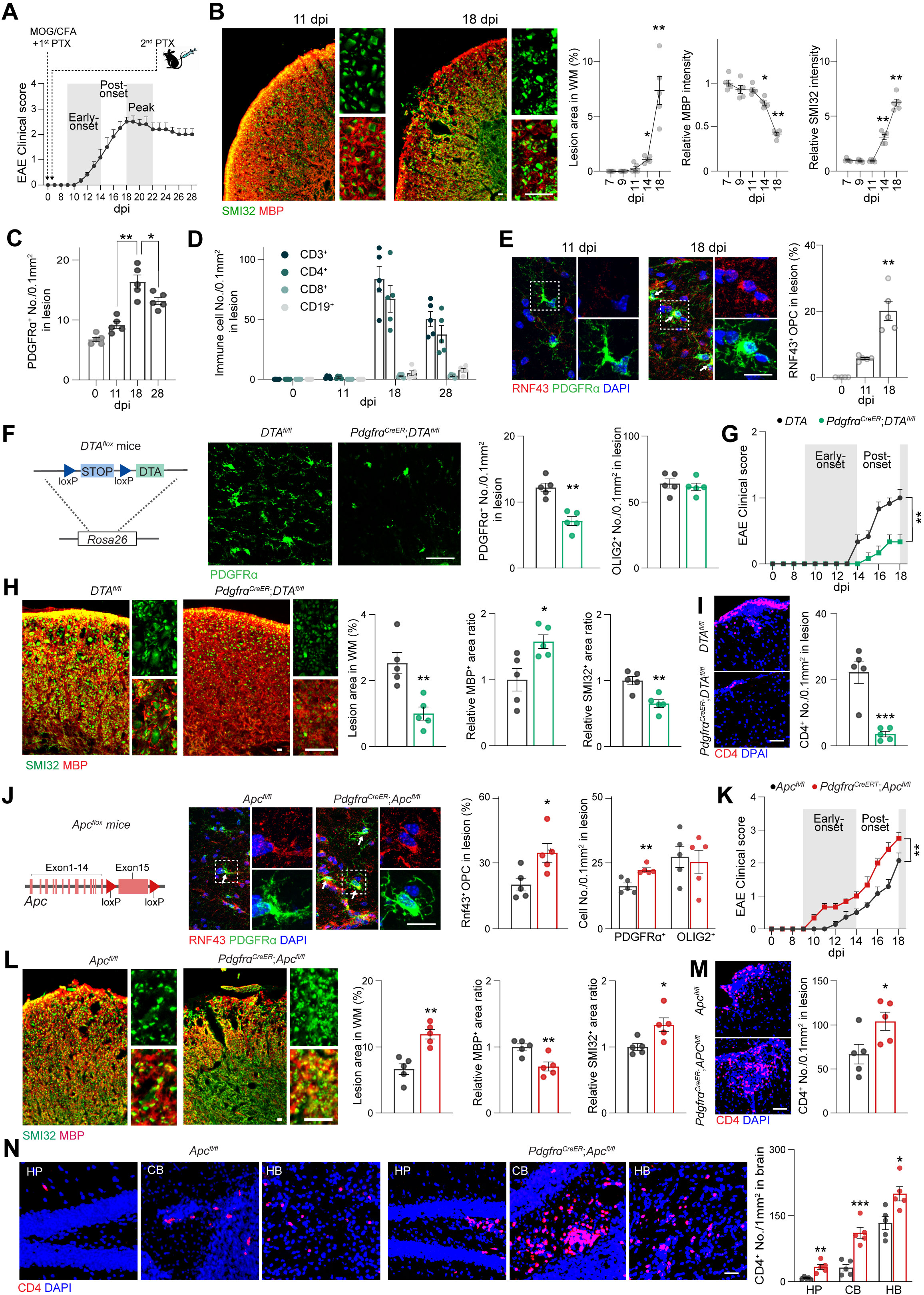
hWnt-OPCs accelerate and exacerbate early EAE and promote CD4^+^ T cell recruitment. (A) Schematic diagram of EAE modeling and mean clinical scores of EAE models over the course of 28 days post-immunization (dpi). C57/BL6 WT mice were immunized using MOG_35-55_ emulsified in CFA by subcutaneous injection on day 0 and received PTX in PBS via intraperitoneal injection on days 0 and 2. (B) Representative immunofluorescent images of the spinal cord show MBP and SMI32 changes at the onset stages of EAE mice. Scale bar represents 20 μm. Quantifications show the changes in relative MBP intensity, relative SMI32 intensity and lesion areas in the white matter (WM) lesions of spinal cords during the EAE progression. (C, D) Quantifications of PDGFRα^+^ OPC cell number and immune cells in WM lesions during the EAE progression. (E) Representative immunofluorescent images and the quantification shows increased RNF43 expression levels in the PDGFRα^+^ OPC during the EAE progression. Arrows highlight the hWnt-OPCs. Scale bar represents 20 μm. (F) Schematic diagram of *DTA^flox^* mice, the representative images of PDGFRα^+^ OPC, and quantifications showing the reduced OPC numbers but unaltered OLIG2+ oligodendroglial lineage cell numbers in *Pdgfr*α*^CreER^*;*DTA^fl/fl^*EAE mice. Scale bar represents 50 μm. (G) Mean clinical scores of *DTA^fl/fl^* and *Pdgfr*α*^CreER^*;*DTA^fl/fl^* EAE mice over the course of 18 dpi. (H) Representative immunofluorescent images of the spinal cord show MBP and SMI32 changes in *DTA* and *Pdgfr*α*^CreER^*;*DTA^fl/fl^*EAE mice at 18 dpi and quantifications of relative MBP intensity, relative SMI32 intensity, and lesion areas. Scale bar represents 20 μm. (I) Representative immunofluorescent images and quantification of accumulated CD4^+^ T cells in lesions of *DTA* and *Pdgfr*α*^CreER^*;*DTA* EAE mice at 18 dpi. Scale bar represents 50 μm. (J) Schematic diagram of *APC^flox^* mice and representative image of RNF43^+^/PDGFRα^+^ OPC. Arrows highlight the activated OPCs. Right panels, quantifications of RNF43+ OPC percentage, as well as PDGFRα^+^ and OLIG2^+^ cell numbers, in *Pdgfr*α*^CreER^*;*Apc^fl/fl^*EAE mice. Scale bar represents 20 μm. (K) Mean clinical scores of *Apc^fl/fl^* and *Pdgfr*α*^CreER^*;*Apc^fl/fl^* EAE mice over the course of 18 dpi. (L) Representative immunofluorescent images of the spinal cord showing MBP and SMI32 changes in *Apc^fl/fl^* and *Pdgfr*α*^CreER^*;*Apc^fl/fl^* EAE mice at 18 dpi and quantifications of relative MBP intensity, relative SMI32 intensity, and lesion areas. Scale bar represents 20 μm. (M) Representative immunofluorescent images and quantification of accumulated CD4^+^ T cells in lesions of *Apc^fl/fl^* and *Pdgfr*α*^CreER^*;*Apc^fl/fl^*EAE mice. Scale bar represents 50 μm. (N) Representative immunofluorescent images and quantification of CD4^+^ T cells in different brain regions of *Apc^fl/fl^* and *Pdgfr*α*^CreER^*;*Apc^fl/fl^* EAE mice. HP: hippocampus, CB: cerebellum, HB: hindbrain. Scale bar represents 50 μm. Plots show Mean ± SEM and \**p* < 0.05, \*\**p* < 0.01, \*\*\**p* < 0.001, n=5 mice/group except specifically mentioned. See also Figure S1.

Following this, we employed a mouse model in which OPCs are eliminated (*Pdgfra^CreER^*;*DTA^fl/fl^*) ^22^ **(Figure 1F)**. It appeared that in OPC-deficient mice, EAE clinical scores were significantly reduced in the post-onset stage **(Figure 1G)**. The severity of demyelination and axonal injury were also alleviated by OPCs elimination **(Figure 1H)**, whereas the density of CD4^+^ T cells decreased concomitantly **(Figure 1I, S1C)**. Thus, a correlation between OPC and EAE progression was established.

OPCs in human MS brains exhibit significantly elevated activation of the Wnt pathway in comparison to their counterparts in current mouse models of MS ^11,12^. In order to assess the role of overactivated Wnt signaling in OPCs which is characteristic for human MS tissues, we utilized another inducible Wnt-activation mouse strain (*Pdgfra^CreER^*;*Apc^fl/fl^*) to trigger overactivation of Wnt in OPCs prior to immunization ^11,12^ **(Figure 1J)**. Such Wnt-overactivation priming of OPCs advanced EAE during the early-onset stage, and exacerbated EAE progression in the post-onset stage **(Figure 1K)**. In line with this, enhanced demyelination and axonal injury were detected in EAE mice with Wnt-overactivated OPCs **(Figure 1L)**, whereas more CD4^+^ T cells were observed in demyelinating lesions **(Figure 1M, S1D)**. Furthermore, accumulated CD4^+^ T cells were found in multiple brain regions of EAE mice with Wnt-overactivated OPCs. In particular, CD4^+^ T cells tended to accumulate in the hippocampus, cerebellum and medulla oblongata **(Figure 1N, S1E)**. Thus, Wnt-overactivated OPCs accelerate pathogenesis of EAE, and prompt CD4^+^ T cell infiltration in its early stage.

### hWnt-OPC-derived CCL4 mediates the CD4^+^ T cell recruitment and EAE pathogenesis

To identify how the Wnt-overactivated OPCs drive immune cell infiltration and EAE pathogenesis, we assessed the migration of MOG_35-55_ activated splenocytes, which were isolated from 2D2 TCR (TCR^MOG^) mice ^23^. Exposure of these splenocytes to conditioned media from Wnt-overactivated OPCs (hWnt-OPC CM), triggered chemotactic response and migration of splenocytes to the lower chamber; whereas treatment with media from healthy OPCs (healthy OPC CM) was ineffective **(Figure S2A)**, thus suggesting that Wnt-overactivated OPCs recruit infiltration of immune cells into the CNS by secreting chemotactic factor(s).

RNA profiling of purified OPCs from the *Pdgfra^CreER^*;*Apc^fl/fl^* mice was compared with OPCs from that non-cre *Apc^fl/fl^*control mice. The gene ontology analysis revealed significant alteration of the Wnt signaling pathway through increasing the gene expression of *Notum*, *Barx1* and *Cdh3* **(Figure 2A)**. The GO analysis also revealed genes responsible for regulation of the immune response, particularly those induced innate immune response, such as *Ccl3, Ccl4, Rgs1*, and *Bcl2a1b*. This was paralleled with an activation of immune response lymphocyte pathways, such as *Ly9, Tlr4, Lilr4b* and *Mcoln2* ^24-26^. Other immune response genes, such as *Lyz1, Lyz2, Cdh3* and *Irf5* were also increased ^27,28^ **(Figure 2A)**. In the volcano plot of upregulated genes, *Ccl4* was highlighted as the most upregulated secreted factor in purified OPCs, along with Wnt pathway indicators, such as *Notum* and *Rnf43*, and other known immunomodulatory factors, such as *Mmp12, Ly9, Lilrb4b* and *Cxcr4* **(Figure 2B)**. Re-analyzing previously published human MS snRNA-seq database confirmed these observations ^29^. Unsupervised clustering of all nuclei into major CNS cell types including neurons, astrocytes, microglia, endothelial cells, oligodendrocytes, OPCs, lymphocytes and pericytes, the OPC cluster was annotated based on *PDGFRα, BCAN, SOX6* and *OLIG2* expressions **(Figure 2C)**. We subsequently evaluated immune cell migration related chemotactic genes (*CCL2, CCL3, CCL4, CCL5 CXCL2, CXCL13, CXCL16*) and their associated receptors (*CCR1, CCR2, CXCR4, CX3CR1*), of which *CCL4/CCL5* were found to be highly expressed by OPCs in MS plaques **(Figure 2D)**. This finding was further corroborated by the markedly increased mRNA expression of *Ccl4* in hWnt-OPCs **(Figure 2E)**, and gradually increased *Ccl4* mRNA levels in the spinal cord during EAE progression **(Figure 2F)**, suggesting that *Ccl4* could be a Wnt targeted downstream gene in OPCs. The disease-specific expression of CCL4 in hWnt-OPCs was also corroborated by immunofluorescent staining. CCL4 was not expressed by OLIG2^+^ oligodendroglial lineage cells in un-immunized tissues, but was excessively expressed in PDGFRα^+^ OPCs at 18 dpi of EAE; it was however rarely detected in CC1^+^ mature oligodendrocytes, microglia, and astrocytes **(Figure 2G, S2B)**, while was not found in PDGFRα^+^ OPCs during neurodevelopmental process **(Figure S2C)**.

**Figure 2.**
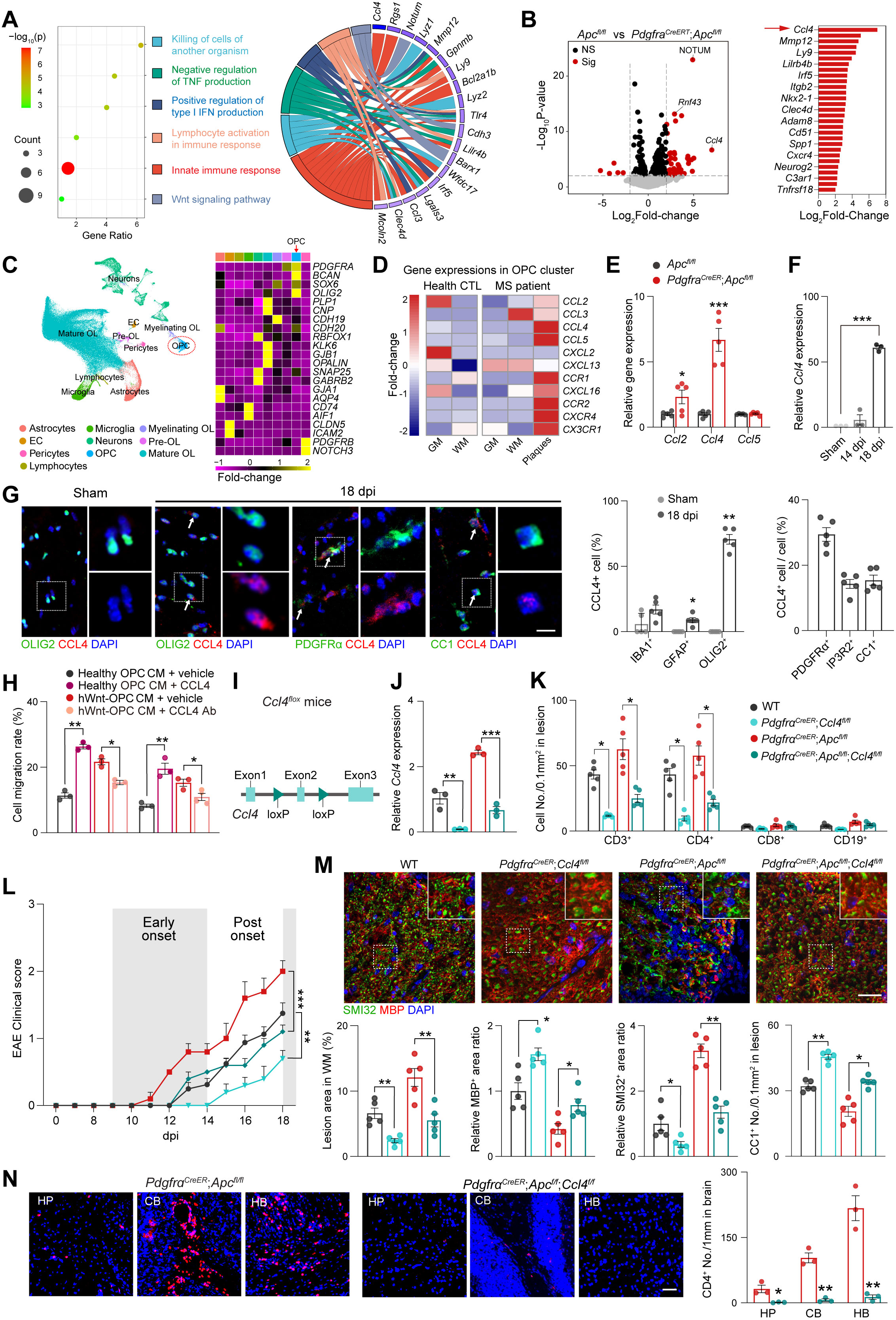
hWnt-OPC-derived CCL4 mediates the CD4^+^ T cell recruitment and EAE pathogenesis. (A) Gene ontology analysis of differentially expressed genes in OPCs from *Apc^fl/fl^* and *Pdgfr*α*^CreER^*;*Apc^fl/fl^*mice. Right panel, Chord diagram displaying genes in the top enriched pathways. (B) Volcano plot of upregulated genes in *Pdgfr*α*^CreER^*;*Apc^fl/fl^* OPCs. Right panel, fold change of top-upregulated gene. (C) UMAP visualization and classification of CNS cellular subclusters from published scRNA-seq database of MS patients. (D) Expressions of chemotactic gene and receptor in hWnt-OPCs deriving from the plaques vs. adjacent white matter of MS patients. (E) qPCR results show the expressions of *Ccl2*, *Ccl4* and *Ccl5* in *Pdgfr*α*^CreER^*;*Apc^fl/fl^* OPCs. (F) qPCR results show the expressions of *Ccl4* changes during the EAE progression. (G) Representative immunofluorescent images of CCL4 and co-staining with oligodendroglial stage markers. Right panels, the proportion of CCL4^+^ cells in glial cell subtypes and oligodendroglial lineage cells in different stages. Scale bar represents 10 μm. (H) Cell migration rates of different types of MOG_35-55_ stimulated immune cells when treated with different conditions. (I) Schematic diagram of *Ccl4^flox^* mice. (J) qPCR results show the expressions of *Ccl4* expression in the OPC with different genotype. (K) Quantification of accumulated CD3^+^ CD4^+^ CD8^+^ and CD19^+^ T cell numbers. (L) Mean clinical scores of *Apc^fl/fl^* and *Pdgfr*α*^CreER^*;*Apc^fl/fl^* EAE mice with or without *Ccl4*-knockout over the course of 18 dpi. (M) Representative immunofluorescent images of the spinal cord show MBP and SMI32 changes in *Apc^fl/fl^* and *Pdgfr*α*^CreER^*;*Apc^fl/fl^* EAE mice with or without *Ccl4*-knockout at 18dpi. Quantification of the percentage of lesion areas, relative MBP intensity, relative SMI32 intensity, and accumulated CC1^+^ cell number. Scale bar represents 50 μm. (N) Representative immunofluorescent images and quantification of CD4^+^ T cells in different brain regions of *Pdgfr*α*^CreER^*;*Apc^fl/f^*^l^ and *Pdgfr*α*^CreER^*;*Apc^fl/fl^*;*Ccl4^fl/fl^* EAE mice. HP: hippocampus, CB: cerebellum, HB: hindbrain. Plots show Mean ± SEM and \**p* < 0.05, \*\**p* < 0.01, \*\*\**p* < 0.01, n=5 mice/group otherwise specifically mentioned. See also Figure S2.

To determine whether the hWnt-OPC specific CCL4 is a trigger for immune cells infiltration, we assessed the effect of CCL4 on migration of activated splenocytes and CD4^+^ T cells isolated from spinal cords of EAE mice. Immune cell migration in healthy OPC CM supplemented with CCL4 ligand increased significantly, while chemotactic effect of hWnt-OPC CM was eliminated by CCL4-blocking antibody **(Figure 2H).** Thus, immune cell infiltration is regulated by CCL4 derived from hWnt-OPCs.

Finally, to examine whether CCL4-mediated immune cell infiltration modifies early EAE *in vivo*, we generated a *Ccl4^fl/fl^* mouse strain and crossed it with the OPC-specific *Pdgfra^CreER^*mice to obtain conditional *Ccl4*-knockout mice which we subjected to EAE protocol **(Figure 2I)**. qPCR showed that the *Ccl4* mRNA levels were significantly reduced by *Ccl4*-conditional knockout **(Figure 2J)**. Consequently, the number of infiltrated immune cells, and CD4^+^ T cells in particular, decreased at 18 dpi, the post-onset stage of EAE **(Figure 2K, S2D)**, confirming that OPC-derived CCL4 plays a dominant role in regulating chemotactic immune cell infiltration into the CNS. Progression of EAE was delayed by *Ccl4*-knockout in both wildtype and OPC Wnt-overactivated mice; while EAE severity was reduced as judged by the declining clinical scores **(Figure 2L)**, reflecting decreased demyelination, axonal injury and increased CC1^+^ mature oligodendrocytes **(Figure 2M, S2D)**. Genetic deletion of CCL4 in OPCs also prevented the infiltration of immune cells into the brain, as shown by the diminished numbers of CD4^+^ T cells in several brain regions **(Figure 2N, S2E)**. However, hWnt-OPCs seem to affect neither proliferation of CD4^+^ T cells **(Figure S3A-C)**, nor activation/differentiation of CD4^+^ T cells **(Figure S3D, E)**. Thus hWnt-OPC-derived CCL4 recruits CD4^+^ T cells into the CNS during early EAE contributing to EAE pathogenesis.

### hWnt-OPCs recruited Th1 cells indirectly instigate demyelination

To determine which subset(s) of CD4^+^ T cells is(are) affected by hWnt-OPCs, we analyzed the proportion of CD4^+^ T cell subsets in the EAE spinal cord. The flow cytometry analysis results revealed that CD4^+^ T cells were mostly represented by T-helper 1 (Th1) cells with Th17 or Treg cells being in minority **(Figure 3A)**. Similarly, the histological analysis confirmed that T-bet^+^ Th1 cells formed a dominant subset of CD4^+^ T cells at 18 dpi **(Figure 3B)**. In addition, more Th1 cells, but not Th17 cells, were observed in EAE with hWnt-OPCs, when compared to wildtype EAE **(Figure 3C, S4A)**. Of note, accumulated CD4^+^ T cells were mainly represented by Th1 subset in the brains of Wnt-overactivated mice **(Figure 3D)**. *In vitro*, by identifying the subsets of translocated CD4^+^ T cells in the lower chamber of a transwell experiment, Th1 cells but not Th17 or Th2 cells, were attracted by hWnt-OPC CM and migrated into the lower chamber **(Figure 3E)**; the migration of Th1 cells in healthy OPC CM supplemented with CCL4 ligand and in hWnt-OPC CM significantly increased, while this chemotactic effect was eliminated in hWnt-OPC CM supplemented with CCL4-blocking antibody **(Figure 3F)**. Following the CCL4-producing OPC population declined at 28 dpi, the late stage of EAE (previously showed in Figure 1C), this proportion of CD4^+^ T cells was altered, as the Th17/Th1 ratio increased at 28 dpi when compared to that of 18 dpi **(Figure S4B)**. Taking into consideration that CCL4 specific receptor, CCR5, is highly expressed by Th1 cells ^30^, our findings indicate that hWnt-OPCs drives a Th1-domainated immune cell infiltration in the early EAE.

**Figure 3.**
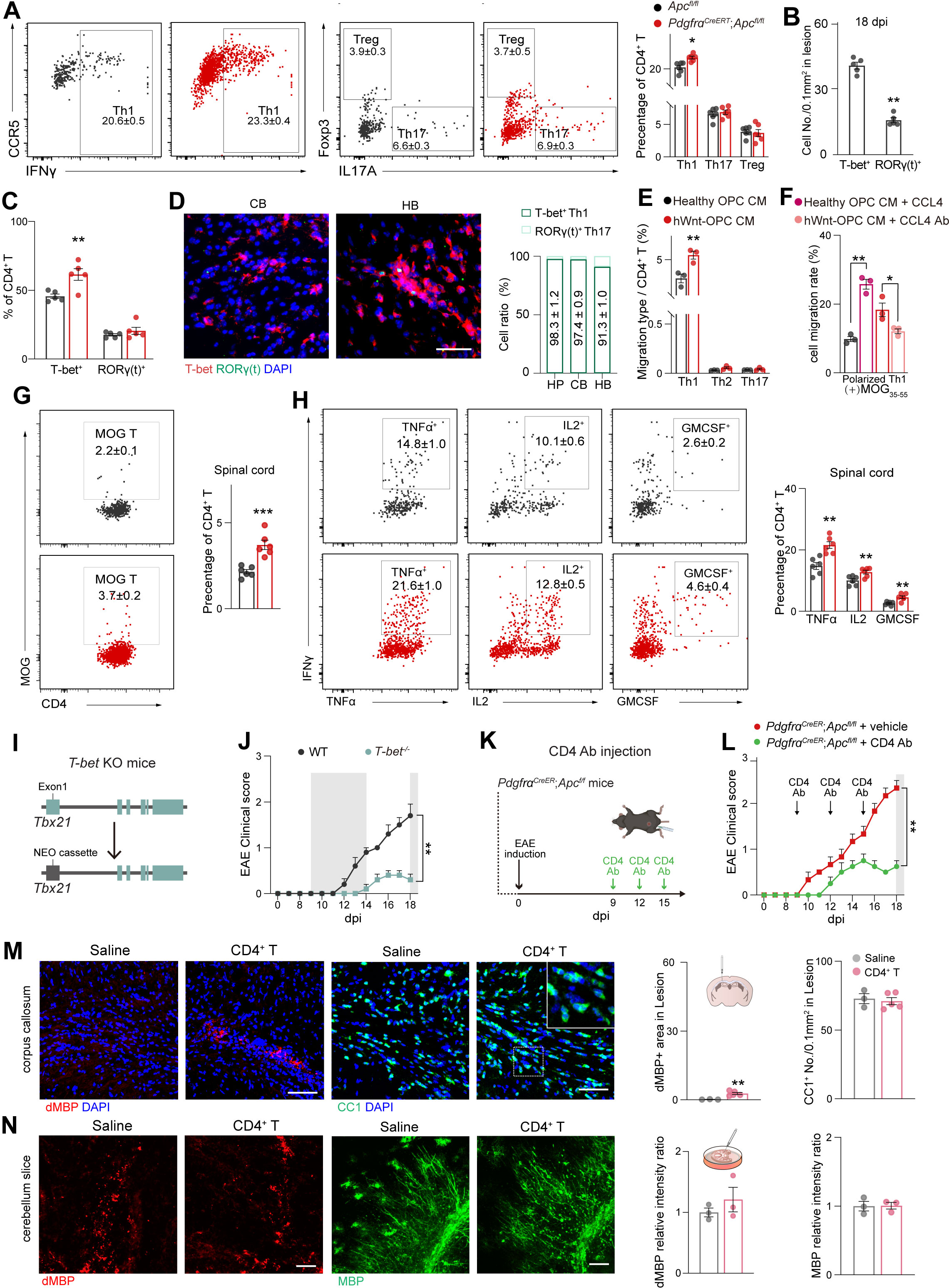
hWnt-OPCs recruited Th1 cells indirectly instigate demyelination. (A) Flow cytometry plots show IFNγ^+^ Th1, IL17A^+^ Th17, and Foxp3^+^ Treg cells in the spinal cord of *Apc^fl/fl^* and *Pdgfr*α*^CreER^*;*Apc^fl/fl^* EAE mice at 18 dpi. (B) Quantification of the proximity of CD4^+^ T cell to OPC in *Apc^fl/fl^* and *Pdgfr*α*^CreER^*;*Apc^fl/fl^* EAE mice at 18dpi. (C) Quantification of the percentage of T-bet^+^ and RORγ(t)^+^ CD4^+^ T cells in *Apc^fl/fl^* and *Pdgfr*α*^CreER^*;*Apc^fl/fl^* EAE mice at 18dpi. (D) Representative immunofluorescent images show Th1 subset cells in different brain regions of *Pdgfr*α*^CreER^*;*Apc^fl/fl^* EAE mice. HP: hippocampus, CB: cerebellum, HB: hindbrain. Scale bar represents 50 μm (E) Flow cytometric quantification of CD4^+^ T cell subsets in the lower chamber from the transwell assay following exposure to healthy OPC CM or hWnt-OPC CM. (F) Cell migration of MOG_35-55_ polarized Th1 cell when treated with healthy OPC CM, healthy OPC CM with CCL4 ligand, hWnt-OPC CM, or hWnt-OPC CM with CCL4-blocking antibody. (G) Flow cytometry shows the frequency of MOG_35-55_ specific CD4^+^ T cells in the spinal cord of *Apc^fl/fl^* and *Pdgfr*α*^CreER^*;*Apc^fl/fl^*EAE mice at 18dpi by executing the MOG_35–55_ tetramer staining for single-cell suspensions from spinal cords of EAE mice. (H) Flow cytometric analysis of TNFα^+^, IL2^+^, and GM-CSF^+^ production in Th1 cells from the spinal cord of *Apc^fl/fl^* and *Pdgfr*α*^CreER^*;*Apc^fl/fl^* EAE mice at 18 dpi. (I) Schematic diagram of *T-bet^-/-^* mice. (J) Mean clinical scores of WT and *T-bet^-/-^* mice over the EAE course of 18 dpi. (K) Schematic diagram of CD4 depletion antibody intraperitoneal injections in *Pdgfr*α*^CreER^*;*Apc^fl/fl^* EAE mice. (L) Mean clinical scores of *Pdgfr*α*^CreER^*;*Apc^fl/fl^* EAE mice with or without CD4 depletion antibody injection. (M) Representative immunofluorescent images and quantification of dMBP and CC1 show mild myelin damages and remained mature oligodendrocytes after saline and CD4^+^ T microinjections. Scale bar represents 100 μm. (N) Representative immunofluorescent images and the quantification of dMBP and MBP show similar myelin damages and the remained myelin-like structures in cultured cerebellum slice after saline and CD4^+^ T microinjections. Scale bar represents 100 μm. Plots show Mean ± SEM and \**p* < 0.05, \*\**p* < 0.01, \*\*\**p* < 0.01, n=5 mice/group otherwise specifically mentioned. See also Figure S3

Next, we tested whether hWnt-OPCs affect the function of infiltrated Th1 cells. The flow cytometry analysis revealed that a significant increase of MOG tetramer-positive CD4^+^ T cells appeared in the EAE mice with hWnt-OPCs, suggesting that hWnt-OPCs attract myelin antigen-specific T cells (MOG tetramer–reactive) **(Figure 3G)**. Furthermore, the percentages of functional cytokine TNF-α, IL-2 and GM-CSF-producing Th1 cells were significantly higher in hWnt-OPC mice **(Figure 3H)**. In addition, the number and proportion of CD69^hi^ CD4^+^ T cell was higher in hWnt-OPC mice. In accordance, CD44^+^CD62L^low^ effector memory CD4^+^ T cells and CD44^+^CD62L^hi^ central memory CD4^+^ T cells were increased in hWnt-OPC mice **(Figure S4C)**. These findings suggest that hWnt-OPCs recruit a greater number of functionally activated Th1 cells in the early EAE, which may accelerate demyelination and axonal damage.

To examine the contribution of infiltrated Th1 to pathophysiology of EAE, we employed the *T-bet^-/-^* mouse strain ^31^, with disrupted expression of T-bet, a transcription factor required for Th1-effector function **(Figure 3I)**. The EAE induced in *T-bet^-/-^* mouse was characterized by delayed progression **(Figure 3J)**. We also modulated the Th1-dominated CD4^+^ T cell population using intraperitoneal injections of CD4 depleting antibody ^32^ in *Pdgfra^CreER^*;*Apc^fl/fl^*EAE mice **(Figure 3K)**. Again, EAE severity was reduced in the early EAE with a higher Wnt background tone **(Figure 3L)**. In the opposite way, we microinjected acutely isolated CD4^+^ T cells from 18 dpi EAE spinal cords into the *in vivo* corpus callosum or in *ex vivo* cerebellar slice cultures, intending to cause demyelination. Unexpectedly, after 3 days of injection, only rather mild immunoreactivity of damaged MBP (dMBP) in both models was detected **(Figure 3M)**. Similarly, the number of CC1-labeled mature oligodendrocytes did not change *in vivo* and the relatively intact MBP-stained myelin-like structures persisted in the *ex vivo* preparation **(Figure 3N)**. Taken together, these findings imply that Th1 cells may not be the direct instigators of fast tissue damages during the early stages of EAE; another cell type is involved.

### A new subpopulation of macrophage-derived cells causes early demyelination

To identify such cell type, we performed randomly down sample from high-dimensional flow cytometry analysis of immune cells isolated from spinal cord tissues in both WT mice and OPC Wnt-overactivated mice at 18 dpi of EAE. Uniform Manifold Approximation and Projection (UMAP) algorithm was used for dimensionality reduction to visualize the data. As a result, a cell cluster co-expressing pattern of NK1.1^+^, CD11b^+^ and Gr-1^+^, was found significantly increased in hWnt-OPC mice **(Figure 4A)**. We further confirmed the presence of this NK1.1^+^CD11b^+^Gr-1^+^ subpopulation at 18 dpi of EAE, in particular, in the EAE spinal cord with hWnt-OPCs **(Figure 4B, C)**. In addition, the density of NK1.1^+^CD11b^+^Gr-1^+^ cell positively correlated with that of infiltrated CD4^+^ T cell and, possibly, with the severity of demyelination in EAE. The density of NK1.1^+^CD11b^+^Gr-1^+^ cells boosted with the increased number of CD4^+^ T cells and exacerbated demyelination in EAE mice with hWnt-OPCs, while NK1.1^+^CD11b^+^Gr-1^+^ cells decreased with the reduced CD4^+^ T cells and prevented demyelination in EAE mice with *Ccl4*-knockout OPCs **(Figure 4D)**, thus suggesting that NK1.1^+^CD11b^+^Gr-1^+^ cells care the likeliest instigators of the early tissue damage.

**Figure 4.**
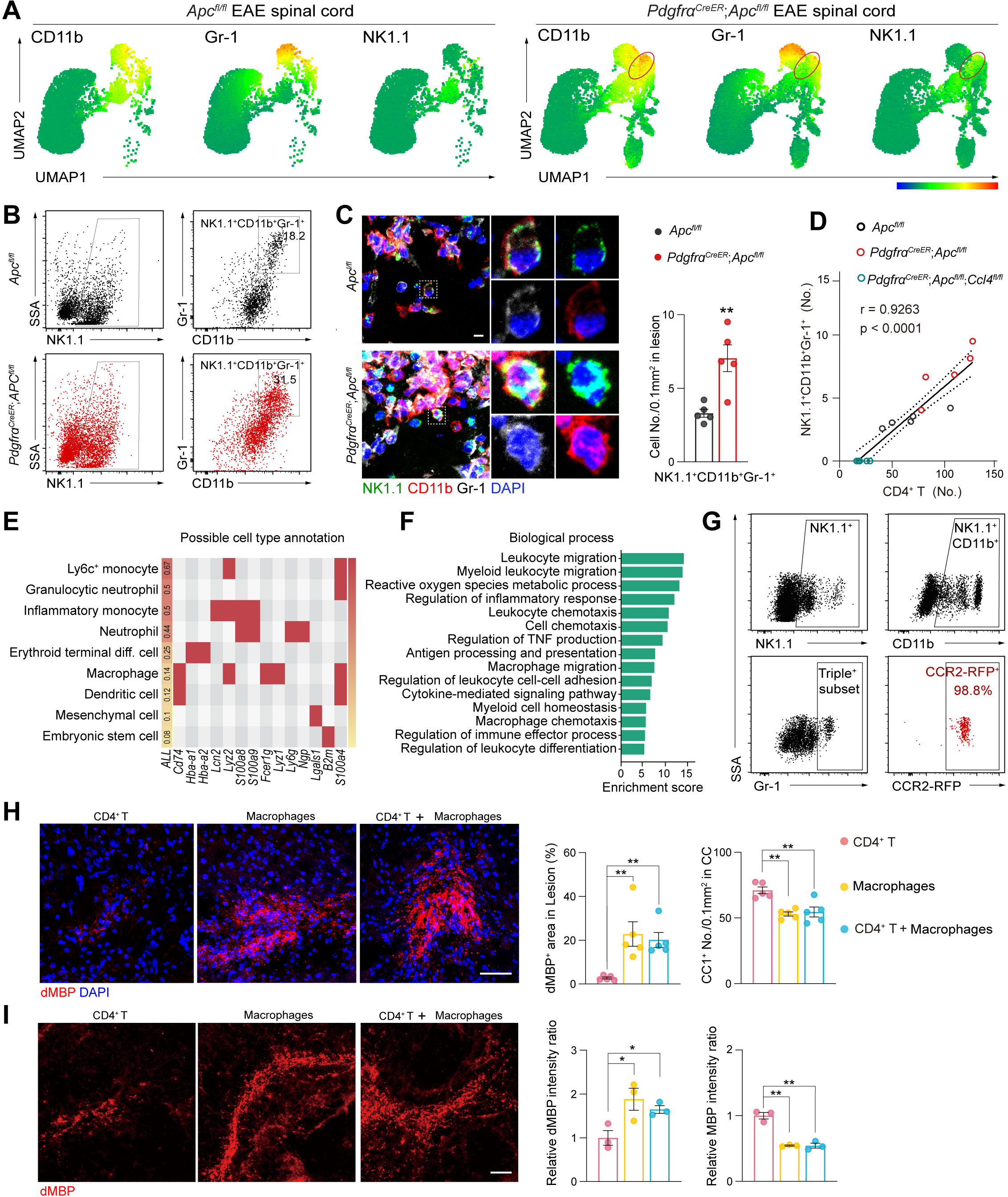
A new subpopulation of macrophage-derived cells causes early demyelination. (A) UMAP visualization of the NK1.1^+^ CD11b^+^ Gr-1^+^ subpopulation in the spinal cord of *Apc^fl/fl^*and *Pdgfr*α*^CreER^*;*Apc^fl/fl^* EAE mice at 18 dpi. (B) Flow cytometry gating reveals NK1.1^+^ CD11b^+^ Gr-1^+^ subpopulation in *Apc^fl/fl^* and *Pdgfr*α*^CreER^*;*Apc^fl/fl^* EAE mice at 18 dpi. (C) Immunofluorescent images confirm the appearance of NK1.1^+^CD11b^+^Gr-1^+^ population in EAE mice spinal cord at 18 dpi. Scale bar represents 10 μm. (D) NK1.1^+^CD11b^+^Gr-1^+^ subpopulation numbers positively correlate with CD4^+^ T cell numbers. Solid and dashed lines represent the simple linear regression and the 95% confidence intervals. (E) The predicted possible cell type annotation by CellMarker 2.0 with highly expressed genes characterized by RNA-seq of NK1.1^+^CD11b^+^Gr-1^+^ cells. (F) Gene ontology analysis of the screened highly expressed genes in NK1.1^+^CD11b^+^Gr-1^+^ cells. (G) Flow cytometric analysis of NK1.1^+^CD11b^+^Gr-1^+^ subpopulation in spinal cord *Olig2^Cre^*;*Apc^fl/fl^*;*Cx3cr1-Gfp*;*Ccr2-Rfp* EAE mice. (H, I) Representative immunofluorescent images of dMBP and quantifications show the demyelination in the corpus callosum and cultured cerebellum slice after cellular microinjections. Scale bar represents 100 μm. Plots show Mean ± SEM and \**p* < 0.05, \*\**p* < 0.01, n=3, 5 mice or independent experiments/group. See also Figure S4 and Figure S5.

To identify the origin of this newly emerged cell subpopulation, RNA-sequencing of acutely isolated NK1.1^+^CD11b^+^Gr-1^+^ cells was performed, and highly expressed genes were processed for cell type annotation ^33^. It appears that the NK1.1^+^CD11b^+^Gr-1^+^ cells are likely a subset of monocytes or macrophages **(Figure 4E)**. We excluded the housekeeping genes and further processed the gene list for pathway and biological process enrichment analysis (**Figure S5A**). The representative enriched terms demonstrated that the highly expressed genes were associated with macrophage migration/chemotaxis, chemokine-related activities, and inflammatory response **(Figure 4F, S5B)**. To acquire a more targeted tracing of the origin of this cell subpopulation, we employed the *Cx3cr1-Gfp*;*Ccr2-Rfp* dual-reporter mouse strain, which allows specific labeling of microglia (by GFP fluorescence) and distinguishing infiltrated macrophages by RFP fluorescence ^34^. The flow cytometry analysis showed that 98.8% of NK1.1^+^CD11b^+^Gr-1^+^ cells were CCR2-RFP positive **(Figure 4G)**. This result was also confirmed by co-staining of NK1.1^+^CD11b^+^Gr-1^+^ cells with RFP by triple-immunofluorescent staining **(Figure S5C)**. Notably, the NK1.1^+^CD11b^+^Gr-1^+^ cells were barely present in the spleen, suggesting that the transformation of this NK1.1^+^CD11b^+^Gr-1^+^ cell subpopulation takes place in the parenchyma rather than in the circulation **(Figure S5D)**, indicating that this newly emerged cell subpopulation are peripheral macrophages that transform into a specific subset under the local inflammatory environment.

To determine whether this subset of macrophage-derived cells mediate rapid demyelination in the early course of EAE, acutely isolated CD4^+^ T cells or macrophage-derived cells were microinjected into the corpus callosum of mouse brains or into cultured cerebellar slices. The dMBP staining was substantially increased in the macrophage-derived cell injection group, and these dMBP levels were comparable between the macrophage-derived cell injection group and the combined group (which has only half number of macrophage-derived cells) **(Figure 4H, I, S5E)**, suggesting that CD4^+^ T cells enhance the capacity of macrophage-derived cells to mediate faster demyelination. This accelerated demyelination was also reflected by other markers, such as declined CC1^+^ mature oligodendrocyte numbers and reduced MBP levels **(Figure S5F, S5G)**. Thus macrophage-derived cells are the instigators that trigger early demyelination in EAE.

### The transformation of macrophage-derived cells is induced by hWnt-OPCs and Th1 cells

We further analyzed the transcription profiles of hWnt-OPCs, CD4^+^ Th1 cells, and macrophage-derived cells to predict the possible interactions ^35^. Results showed that the network was enriched in the chemotaxis biological process between macrophage-derived cells and Th1 cells, while enriched in the inflammatory response biological process between macrophage-derived cells and hWnt-OPCs **(Figure S6A)**.

Given Th1 cells have a central role in macrophage activation ^36^, we firstly determined the regulatory effects of Th1 cells on macrophage-derived cells. We employed the *T-bet^-/-^* mouse strain ^31^, in which the functional CD4^+^ T cells are markedly reduced. In the early stage of EAE within *T-bet^-/-^* model, the number of OPCs was not altered but the numbers of CD4^+^ T cells and macrophage-derived cells declined concomitantly **(Figure 5A, B, C, S6B)**. We also modulated CD4^+^ T cells using intraperitoneal injections of CD4 depleting antibody ^32^ in *Pdgfra^CreER^*;*Apc^fl/fl^*EAE mice. Again, the number of OPCs did not change, whereas number of CD4^+^ T cells and macrophage-derived cells were significantly decreased **(Figure 5D, E, F, S6C)**. These translated in significantly lower level of demyelination and axonal injury as compared to wildtype or control EAE mice **(Figure 5G, H),** and reduced EAE severity (Figure 3J, L).

**Figure 5.**
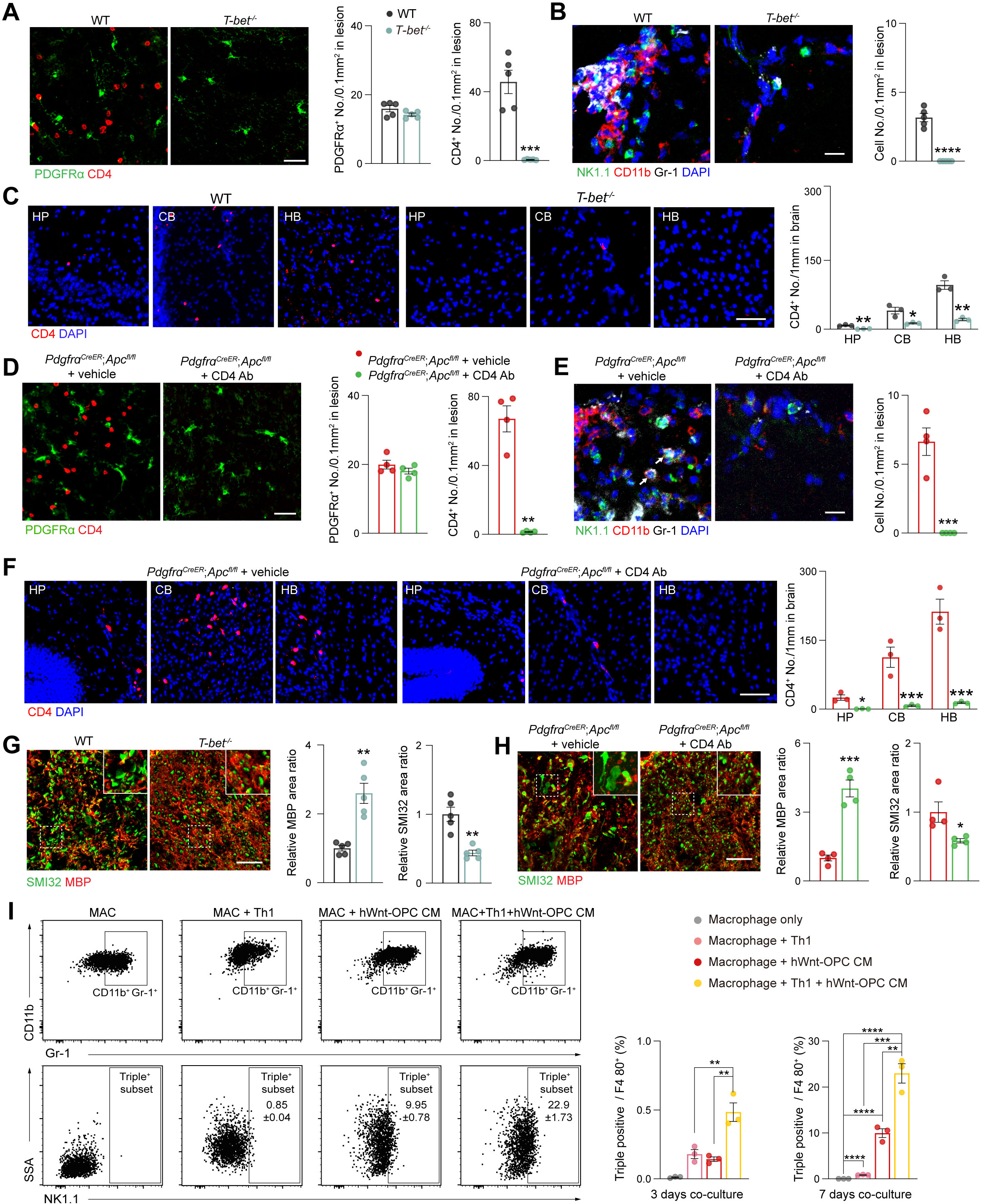
The transformation of macrophage-derived cells is induced by Th1 cells and hWnt-OPCs. (A) Representative image and quantifications show the number of OPCs are not altered but the number of CD4^+^ T cells decline significantly in *T-bet^-/-^* EAE mice. (B) Representative image and quantification of macrophage-derived cells in lesions of WT and *T-bet^-/-^* EAE mice. Scale bar represents 20 μm. (C) Representative immunofluorescent images and quantification of CD4^+^ T cells in different brain regions of WT and *T-bet^-/-^* EAE mice. Scale bar represents 50 μm. (D) Representative image and quantifications show the number of OPCs are not altered but the number of CD4^+^ T cells decline significantly in CD4 depletion antibody injected *Pdgfr*α*^CreER^*;*Apc^fl/fl^* EAE mice. (E) Representative image and quantification of macrophage-derived cells in lesions of *Pdgfr*α*^CreER^*;*Apc^fl/fl^*EAE mice with or without CD4 depletion antibody injection. Scale bar represents 20 μm. (F) Representative immunofluorescent images and quantification of CD4^+^ T cells in different brain regions of *Pdgfr*α*^CreER^*;*Apc^fl/fl^* EAE mice with or without CD4 depletion antibody injection. Scale bar represents 50 μm. (G) Representative immunofluorescent images of the spinal cord and quantification show MBP and SMI32 changes in WT and *T-bet^-/-^* EAE mice at 18 dpi. Scale bar represents 50 μm. (H) Representative immunofluorescent images of the spinal cord and quantifications show MBP and SMI32 changes in *Pdgfr*α*^CreER^*;*Apc^fl/fl^* EAE mice with or without CD4 depletion antibody injection. Scale bar represents 50 μm. (I) Flow cytometric analysis shows 7 days co-culture of primary macrophages and Th1 cells under different conditions. Statistical analysis showed the percentage of triple positive macrophages significantly increased when primary macrophages co-cultured with Th1 cells and WNT-OPC CM for 3 days and 7 days. Plots show Mean ± SEM and \**p* < 0.05, \*\**p* < 0.01, \*\*\**p* < 0.001, n=3, 5 mice or independent experiments/group. See also Figure S6.

Macrophage-derived cells also could be affected by hWnt-OPCs (Figure S6A), therefore we assessed the regulatory effects of Th1 cells and hWnt-OPCs on macrophages. The flow cytometry analysis showed that both Th1 cells and hWnt-OPC CM were able to induce the transformation of peritoneal macrophages from primary state to specific pathological state. Unexpectedly, hWnt-OPC CM had a stronger effect (induced 9.95% triple positive macrophages at 7 days co-culture) when compared to Th1 cells (0.85%). Combination of Th1 cells and hWnt-OPCs most significantly enhanced this transformation by increasing the triple positive subpopulation of macrophages (22.9%) **(Figure 5I, S6D)**, suggesting that hWnt-OPCs and Th1 cells cooperate to exert a stronger effect on macrophage transformation. This result was also confirmed by the triple-immunofluorescent staining of NK1.1^+^CD11b^+^Gr-1^+^ cells in co-cultures **(Figure S6E)**. These findings indicate that the transformation of macrophage-derived cells requires the assistance of both Th1 cells and hWnt-OPCs. Taking into consideration that hWnt-OPCs also recruit Th1 cells in EAE (Figure 3), hWnt-OPCs thus act as a driver for the immune cascade network at the early stage of EAE.

### Macrophage-derived cells are cytotoxic and activate Wnt pathway in OPCs

The gene profiles of macrophage-derived cells further characterized their potential functions, by showing highly expressed cytotoxic factors, such as *S100a8/9, B2m* and *Il18* ^37-39^ **(Figure 6A)**. Gene set enrichment analysis similarly highlighted cell cytotoxicity, apoptosis, necroptosis and pyroptosis related gene sets were enriched by the high expression of the target genes in macrophage-derived cells ^40^ **(Figure 6B)**.

**Figure 6.**
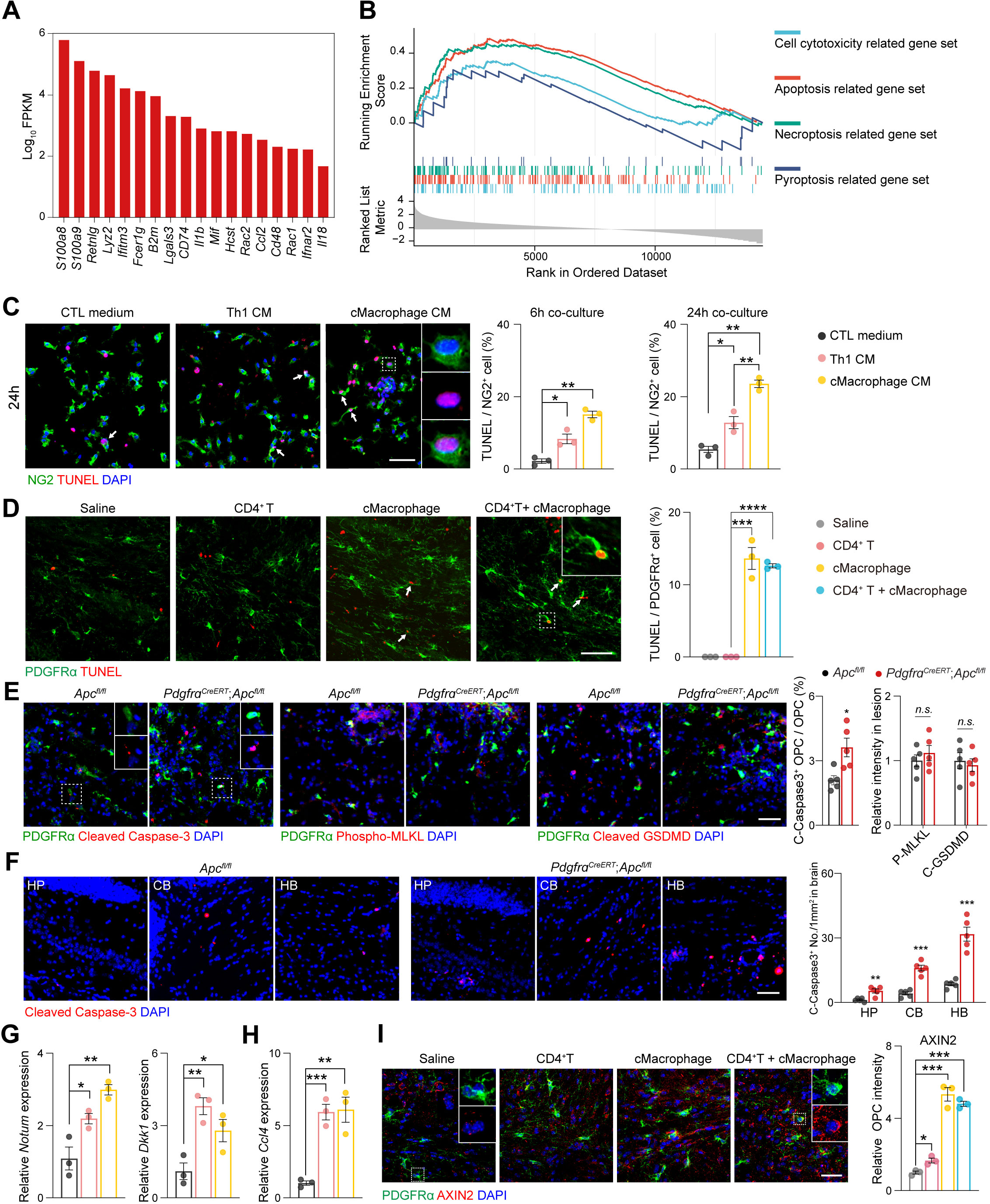
Macrophage-derived cells are cytotoxic and activate Wnt pathway in OPCs. (A, B) Gene set enrichment analysis of cell cytotoxicity, apoptosis, necroptosis and pyroptosis related genes in macrophage-derived cells. (C) Tunel staining in OPCs shows an increased OPC apoptosis rate when exposed to immune cellular conditioned media and with longer exposure time. Scale bar represents 50 μm. (D) Tunel staining in OPCs in the lesions with different immune cell microinjections. Scale bar represents 50 μm. (E) Representative immunofluorescent images and quantification of Cleaved Caspase-3, Phospho-MLKL and Cleaved GSDMD in lesions of *Apc^fl/fl^* and *Pdgfr*α*^CreER^*;*Apc^fl/fl^*EAE mice. Scale bar represents 50 μm. (F) Representative immunofluorescent images and quantification of Cleaved Caspase-3 in different brain regions of *Apc^fl/fl^* and *Pdgfr*α*^CreER^*;*Apc^fl/fl^* EAE mice. HP: hippocampus, CB: cerebellum, HB: hindbrain. Scale bar represents 50 μm. (G, H) Relative mRNA expression of Wnt activation indicators *Notum*, *Dkk1*, and *Ccl4* in OPCs when treated with different conditions. (I) Representative immunofluorescent images and quantification of AXIN2 show an upregulated Wnt pathway activation in OPCs in the lesions with cellular microinjections. Scale bar represents 50 μm. Mean ± SEM and \**p* < 0.05, \*\**p* < 0.01, \*\*\**p* < 0.001, \*\*\*\**p* < 0.0001, n=3, 5 mice or independent experiments/group.

To confirm this cytotoxic effect, cultured OPCs were exposed to the macrophage-derived CM or Th1 CM. More Tunel-positive OPCs were present after treatment with macrophage-derived CM than with Th1 CM; while the OPC apoptosis rate was significantly increased in macrophage-derived CM with longer exposure time **(Figure 6C)**. Similarly, *in vivo* Tunel-positive OPCs were only found in lesion areas with injections of macrophage-derived cells; combining cell types exerted the same effect as macrophage-derived cells alone **(Figure 6D)**, indicating a dominant role of macrophage-derived cells in inducing OPC death. In addition, cleaved Caspase-3 positive but phospho-MLKL (necroptosis) or cleaved GSDMD (pyroptosis) negative OPCs **(Figure 6E)** indicated an apoptosis-mediated OPC death in EAE progression. Strikingly, this cell apoptosis was also detected in several brain regions **(Figure 6F)**, implying an extended EAE pathology in the brain. These results support our previous findings, which showed declined OPC numbers at 28 dpi, the late stage of EAE (Figure 1C), and were consistent with the idea that when immune cells infiltrate the CNS, they induce tissue damage, including OPC death ^1^. Taking into consideration that macrophage-derived cells could express multiple cytotoxic factors and induce cell death of mature oligodendrocytes (Figure 4H, I, S5F), we termed them as cytotoxic macrophages.

Given that cytotoxic macrophage produced TNFα (Figure 4F) ^41^ while Th1 cells ^42^ regulate the Wnt pathway activation, our qPCR results demonstrated that the mRNA levels of Wnt activation indicators *Dkk1* and *Notum* **(Figure 6G)**, and *Ccl4* **(Figure 6H)** were significantly upregulated by both macrophage-derived CM and Th1 CM treatments. Confirmative findings were obtained *in vivo*. Upregulated Wnt-activation indicator AXIN2 was found in OPCs in the lesioned areas after injections of CD4^+^ T cells, but much more significantly increased activation of Wnt pathway was present after the injection of cytotoxic macrophages, and the injection of combined cell types showed the same effect as cytotoxic macrophages alone **(Figure 6I)**. These findings indicate that infiltrated immune cells, in particular cytotoxic macrophages interact with OPCs, upregulating the Wnt activity and CCL4 expression in hWnt-OPCs, further increasing hWnt-OPCs population and creating an OPC-immune cascade, which consequently, induced death of hWnt-OPCs.

### Breaking the OPC-immune cascade by targeting CCL4 attenuates EAE

To explore the therapeutic potential of our findings, we treated WT and *Pdgfra^CreER^*;*Apc^fl/fl^* EAE mice with maraviroc (MVC), a selective antagonist of CCR5^43^, the receptor of CCL4 highly expressed by Th1 cells both in MS and EAE ^44,45^. Treatment with MVC eliminated the chemotactic effect of hWnt-OPC CM on Th1 cells in a transwell experiment **(Figure 7A)**. Clinical scores indicated that EAE progression was delayed by MVC administration during the early-onset stage, and the clinical scores were also improved in the post-onset stage **(Figure 7B)**. Dual labelling of SMI32/MBP at 18 dpi also confirmed that early demyelination was rescued by using MVC to target CCL4 **(Figure 7C)**. The number of CD4^+^ T cells (in particular, the T-bet^+^ Th1 cells) and cytotoxic macrophages counted at 18 dpi was significantly reduced by MVC administration **(Figure 7D, E)**. Beneficial effects of MVC were also seen for the CD3^+^ lymphocytes, CD8^+^ T cells and CD19^+^ B cells **(Figure 7D)**. Furthermore, MVC administration prevented the CD4^+^ T cell infiltration into several brain regions, such as the hippocampus, cerebellum, and medulla **(Figure 7F, S7A)**.

**Figure 7.**
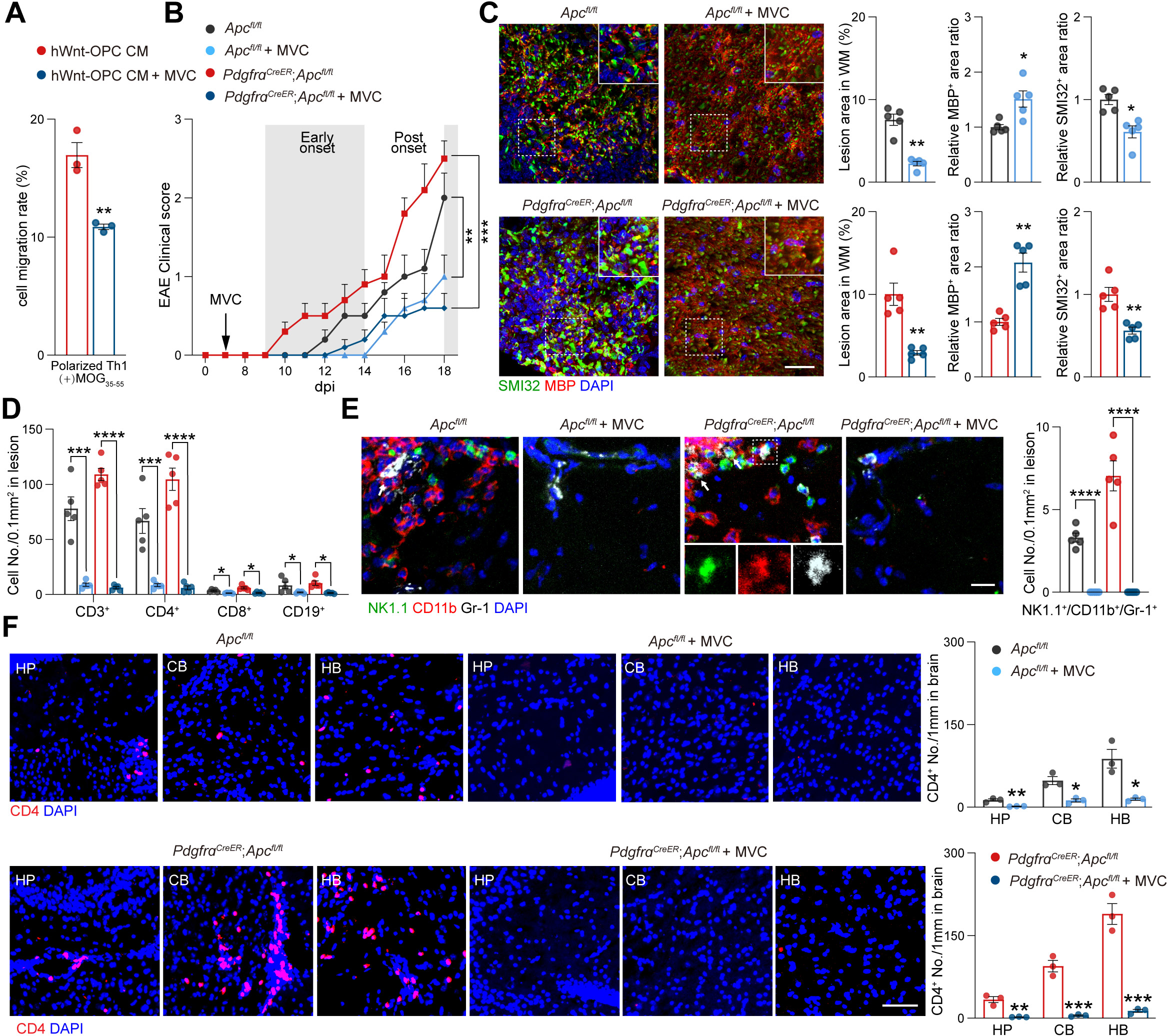
Breaking the OPC-immune cascade by targeting CCL4 attenuates EAE pathologies. (A) Cell migration rate of MOG_35-55_ polarized Th1 cell when treated with different OPC-conditional media. (B) Mean clinical scores of *Apc^fl/fl^* and *Pdgfr*α*^CreER^*;*Apc^fl/fl^*EAE mice with or without MVC treatment over the EAE course of 18 dpi. (C) Representative immunofluorescent images of the spinal cord and quantifications show MBP and SMI32 changes in MVC-treated *Apc^fl/fl^*and *Pdgfr*α*^CreER^*;*Apc^fl/fl^*EAE mice at 18 dpi. Scale bar represents 50 μm. (D) Quantification of infiltrated immune cells in lesions of EAE mice with or without MVC treatments. (E) Immunofluorescent images of NK1.1^+^CD11b^+^Gr-1^+^ cytotoxic macrophages confirm the reduction of this subpopulation in EAE mice after MVC treatments. Scale bar represents 20 μm. (F) Representative immunofluorescent images and quantification of CD4^+^ T cells in different brain regions in *Apc^fl/fl^* and *Pdgfr*α*^CreER^*;*Apc^fl/fl^*EAE mice with or without MVC treatments. Scale bar represents 50 μm. Mean ± SEM and \**p* < 0.05, \*\**p* < 0.01, \*\*\**p* < 0.001, \*\*\*\**p* < 0.0001, n=3, 5 mice/group otherwise specifically mentioned. See also Figure S7.

To exclude the direct action of MVC on OPCs, purified OPCs were treated with MVC in proliferating and differentiating conditions. Neither OPC propagation nor OPC maturation was altered by MVC treatment **(Figure S7B)**. Together, our results indicate that MVC can break the OPC-immune cascade, reduce immune cell infiltration at early stage of EAE, and attenuate EAE progression.

## Discussion

The EAE animal model of MS does not fully recapitulate pathophysiology and progression of the human MS; in particular, EAE lesions are confined to the spinal cord, whereas human disease primarily affects the brain. We argue that the difference is in OPCs, which in the human pathology undergo pathological transformation caused by an overactivation of Wnt signaling cascade. Here we mimicked human pathology by artificially increasing the Wnt tone in mice OPCs, which translated into a much accelerated and exacerbated EAE pathology; moreover, this population of pathological OPCs extended EAE pathology to the brain.

Thus, in the present study, we demonstrate a previously unknown role for pathological hWnt-OPCs as hubs of an immune cascade network defining the early course of EAE progression. These pathological hWnt-OPCs are distinguished by an overactivated Wnt signaling, and expression of CCL4 (which is virtually absent in healthy OPCs), which mediates recruitment of Th1 cells into the CNS. The Th1 cells accumulate at the sites of primary damage and cooperate with hWnt-OPCs to induce the transformation of cytotoxic macrophages that execute the primary damage to the myelin. Infiltrated immune cells in turn interact with hWnt-OPCs thus establishing a vicious cycle of early and progressive tissue damage. These findings have significant implications for our understanding how pathological OPCs together with invading immune cells, including previously unknown cytotoxic macrophages define MS pathophysiology and present a target for therapeutic interventions.

Emerging evidence suggests that OPCs, and especially adult OPCs, have multiple physiological roles and assume complex pathophysiological significance in various neurological diseases ^46^. OPCs are not only responsible for myelin formation in development, or myelin repair in diseases, but they can also regulate angiogenesis in the postnatal developing CNS and are engaged in intricate interactions with astrocytes and neurons in cognitive disorders ^46-49^. Previous studies demonstrated an increase in OPC number in early MS as well as in the early stages of EAE ^16,17,50,51^. Furthermore, the accumulation of OPCs positively correlates with the early infiltration of immune cells ^52,53^. These findings question the OPCs purported role as ‘victims’ of immune attack, but imply an immunomodulatory capability as well as active participation of OPCs in pathophysiology and progression of MS ^13,14,54,55^. Our results demonstrate that hWnt-OPCs trigger inflammatory responses, even in the brains of EAE mice. Consequently, manipulations with pathological hWnt-OPCs may affect, slow down or even arrest the progression of MS.

Activation of Wnt signaling cascade has different pathological roles in MS, which depend on the specific cell types. For instance, activated Wnt pathway in endothelial cells reduces immune cell infiltration ^9^, which may alleviate CNS inflammation. However, Wnt-overactivation in OPCs contributes to the OPC perivascular clustering and disruption of BBB ^11,12^, thus amplifying CNS inflammation. Notably, CCL4, which is the downstream of Wnt pathway in OPCs, is increased in the cerebrospinal fluid of relapsing/remitting MS and in active MS lesion tissues ^56^. Taking into consideration that CCL4 specific receptor, CCR5, is also over-expressed in lesion-derived T cells and peripheral T cells in MS ^44,45^, we demonstrate that pathological OPCs induce Th1 cell infiltration through a Wnt-CCL4 axis. Given that myelin gene MOG-driven specific oligodendrocyte ablation aborts adaptive immune response in the mouse spinal cord ^57^, we suggest that OPCs are more immunomodulatory than mature oligodendrocytes.

Multiple immune cell types have been observed in early MS tissues ^1,58,59^. It was believed that myelin-reactive Th1 cells initiate EAE ^4,5^, however, its role in demyelination was challenged by the discovery of Th17 cells ^6,7^. In the present study, we show that Th1 cells accumulate in the EAE tissues before Th17 cells, CD8^+^ T cells, and CD19^+^ B cells, but do not induce fast demyelination on their own, being in need of cooperation with another effector cells. A newly identified subpopulation of NK1.1^+^CD11b^+^Gr-1^+^ cells (which we dubbed cytotoxic macrophage), originates from macrophage, expresses multiple cytotoxic factors, and is the main executor of tissue damages in early EAE. These cytotoxic macrophages are induced by Th1 cells ^60^ and hWnt-OPCs respectively, while hWnt-OPCs together with Th1 cells exert a much stronger effect on the macrophage transformation, suggesting that Th1 cells play a critical role in promoting this transformation. Although both beneficial and detrimental roles of macrophages were found at different stages of MS and EAE ^15^, analysis of *Cx3cr1-Gfp*;*Ccr2-Rfp* dual-reporter mice supports our findings by showing that CCR2-RFP^+^ macrophages increase at the onset of EAE but decrease after the peak stage ^61^. Such time course is also consistent with the findings in early and active lesions in MS, but is lower in inactive, demyelinated, or remyelinating lesions ^2^. Simultaneously, infiltrated Th1 cells and cytotoxic macrophages in their turn interact with OPCs, stimulating a further Wnt activation, as reported in other cell types ^62,63^. Consequently, the expression of CCL4 further increases, expanding this OPC-immune interaction through positive feedback. Considering that immune cells (including CD4^+^ T cells and CCR2-RFP^+^ macrophages) extravasate into the CNS in development (i.e. in the absence of EAE) when the Wnt pathway is overactivated in OPCs ^11^, it is tempting to speculate that hWnt-OPC-immune cellular network may emerge during the onset of the disease or in a newly forming plague in the surrounding normal-appearing white matter **(Figure S7C)**. Initially a few OPCs interact with extravasated immune cells, and this interaction converts them into hWnt-OPCs. Then, hWnt-OPC-derived CCL4 acts on Th1 cell infiltration, and subsequent macrophage transformation into cytotoxic macrophages. The latter in turn contribute to further increase Wnt activation and CCL4 expression in neighboring OPCs. Ultimately, a vicious cycle of hWnt-OPCs and immune cells emerges, recruiting more immune cells in a broader area, leading to demyelination, axonal degeneration, OPC death and progressive tissue damage. In later stages of MS, following the decrease of OPCs and increase of Th17 cells and B cells ^16,17^, different immunological cells may cause irreversible nerve injuries by different mechanisms^15,58^.

Several therapies ^64-67^, with Rituximab, Inebilizumab, Obinutuzumab, were used to target B cells in neuromyelitis optica spectrum disorder, MOG antibody-associated disease, or relapsing-remitting MS. An early intervention, targeting immune cells to prevent early tissue damage and irreversible neuropathological outcomes, seems to a suitable alternative, yet hitherto missing, strategy. Although CD4^+^ T cell depletion had no clinical benefit ^58,68^, a recent study revealed that by inducing apoptosis of reactive CD4^+^ T cells and immunosuppressive monocytes by targeting oligodendrocyte-released extracellular vesicles the severity of EAE can be reduced ^69^. We, likewise, suggest that an effective early intervention for MS can target upstream regulators of the autoimmune reaction represented by hWnt-OPCs. Given that CCL4 was considered as a biomarker for MS severity ^70^, and it mediates the hWnt-OPCs driven pathological cascade during the onset of EAE, we tested a European Medicines Agency-approved HIV treatment drug MVC with a reported immunotherapeutic effect on EAE ^71,72^. By breaking the CCL4-mediated OPC-immune cascade in the early EAE with MVC, we were not only able to reduce the immune cell infiltration to the spinal cord and the brain, but also to delay EAE pathogenesis and alleviate severity of the tissue damage. It is plausible to suggest that by combining immunomodulatory drugs with agents preventing pathological transformation of OPCs the disease progression may be effectively managed ^12, 73-75^.

In conclusion, our findings suggest that pathological hWnt-OPCs are the initiator of a vicious cycle involving immune cells, which trigger immunomodulatory network, and propose a potential early intervention strategy for MS treatment.

## Limitations of the study

This study was designed to investigate the immunomodulatory network in early EAE, in which accumulated immunomodulatory cells, together with pathological hWnt-OPCs, interact to cause tissue damage. Although we observed more immune cells infiltrating the brains of mice with overactivated Wnt, further studies are required to verify our findings in human MS subjects and larger prospective cohorts. An additional limitation of our study is that we analyzed the OPC-immune cell interactions in fixed tissues and *primary* cultures. In future studies, new generations of multiple fluorescence-labeled reporter mice could help to visualize the OPC-immune cell interactions, especially the OPC-cytotoxic macrophage interaction, and analyze their trajectories in live animals. Despite these limitations, our work highlights a previously unknown pathological hWnt-OPC-initiated immune cascade as a new immunotherapeutic target for early intervention in MS.

## Acknowledgments

This work was supported by grants from the National Nature Science Foundation of China (NSFC 32070964 and 32271034 to J.N.; 81971309 and 32170980 to C.Y.; 82200562 to T.H.). National Key Research and Development Program of China (2021ZD0201703 to J.N.). Chongqing Natural Science Fund for Distinguished Young Scholars (CSTB2023NSC Q-JQX0030 to J.N.). Science Fund for Creative Research Groups of the Natural Science Foundation of Chongqing (cstc2019jcyj-cxttX0005 to Q.Y.). Guangdong Basic and Applied Basic Research Foundation (2022B1515020012 to C.Y.; 2022A1515111165 to Q.W.; 2021A1515110268 and 2023A1515010651 to Y.S.; 2021A1515110121 to T.H.). Shenzhen Fundamental Research Program (JCYJ20210324123212035, RCYX20200714114644167, and ZDSYS20220606100801003 to C.Y.; JCYJ20220530144816038 and RCBS20221008093118042 to Q.W.; RCBS20210706092411028 and JCYJ20210324121214039 to Y.S.; JCYJ20210324122809025 to T.H.).

## Author contributions

J.N., C.Y. and Y.W. conceived the study. J.N., C.Y. and Y.W. designed the experiments. Q.W., T.H., Z.Z., Y.S., Z.W., G.Y., Y.L., X.W., H.L., X.C., Z.J., Q.Y. and J.N. performed the experiments and analyzed the data. Q.W., Y.L., Z.Z. and Y.W. performed analysis of the flow cytometry and RNA-seq data. J.Z., Y.Z., Y.T. and Q.Y. contributed to discussion. A.V. helped design some of the experiments and contributed to discussion. J.N., C.Y. and Y.W. wrote the manuscript. A.V. made the editing.

## Methods

### Mice

All mouse strains were maintained in the Third Military Medical University specific pathogen-free animal facility, and the animal procedures were performed according to the guidelines of laboratory animal welfare and ethics committee of the Third Military Medical University. Mouse ages were identified in the main text and figure legends. Both mouse genders were used in this study. Sex-matched mice were randomly assigned to the different experimental groups. Littermate controls were used where appropriate.

<u>*Pdgfr*α*^CreER^*</u>: this mouse strain has been described previously ^76^. This mouse was used to cross with other floxed mice strains to generate conditional knockout mouse models when tamoxifen was given.

<u>*Olig2^Cre^*</u>: this mouse strain has been described previously ^77^. This mouse strain was used to knockout *Apc* in oligodendroglial lineage cells during EAE progression.

<u>*Apc^flox^*</u>: this strain was crossed with *Olig2^Cre^*or *Pdgfr*α*^CreER^* to over-activate Wnt pathways in OPCs as reported in our previous studies ^11,12,78^.

<u>*DTA^flox^*</u>: this strain was crossed with *Pdgfr*α*^CreER^* to achieve OPC-specific diphtheria toxin expression and subsequent OPC elimination; the non-cre;*DTA* littermates were used as the control mice ^79^.

*<u>2D2:</u>* 2D2 TCR (TCR^MOG^) mouse strain has been described previously ^23^. Splenocytes isolated from this mouse strain express a myelin oligodendrocyte glycoprotein (MOG)-specific T cell receptor (TCR), and respond to MOG_35-55_ peptide stimulation *in vitro*.

<u>*Ccl4^flox^*</u>: this mouse strain was newly generated in this study using the CRISPR-Cas9 technology by insertion of loxP sites flanking exon2 of *Ccl4* to disrupt the function of CCL4 protein. The CRISPR/Cas9 system was microinjected into the fertilized eggs of C57BL/6J mice. Fertilized eggs were transplanted to pseudo-pregnant mice at the blastocyst stage to obtain positive F0 mice, following confirmed by PCR and sequencing. A stable F1 generation mouse model was obtained by mating positive F0 generation mice with C57BL/6 mice. These procedures were performed by Gempharmatech Co., Ltd. in China. We used a breeding strategy of crossing either *Pdgfr*α*^CreER^*;*Ccl4^fl/fl^* mice or *Pdgfr*α*^CreER^*;*Apc^fl/fl^*;*Ccl4^fl/fl^* mice, thus generating OPC-specific *Ccl4* knockout or *Apc*/*Ccl4* double knockout mouse strains.

<u>*Cx3cr1-Gfp*;*Ccr2-Rfp*</u>: this reporter mouse strain specifically labels microglia in the CNS by GFP, and macrophages by RFP ^34^. By using this mouse strain, we can distinguish infiltrating macrophages from resident microglia within lesions ^11^.

<u>*T-bet^-/-^*</u>: this mouse strain has been described previously ^31^, and purchased from the Jackson laboratory. In this mouse strain, the expression of T-bet was disrupted, resulting in a markedly reduced population of functional CD4^+^ T cells.

### EAE mouse model

EAE models were induced using MOG_35-55_ following the previously described protocol ^80^. Briefly, 8-10-week-old mice were subcutaneously injected in the flank with 200 μg MOG_35-55_ in CFA emulsion containing 4 mg/mL of heat-killed Mycobacterium tuberculosis H37Ra at the day of post-immunization (0 dpi). Mice also received 200 ng pertussis toxin (PTX) in PBS via intraperitoneal injection at 0 and 2 dpi. Then, immunized mice were monitored and scored daily for disease signs according to the following 5-point clinical scoring criteria with 0.5-point increments as follows: 0: no disease; 1: full limp tail; 2: severe gait impairment/hindlimb paresis; 3: hindlimb paralysis; 4: hindlimb paralysis and partial forelimb paralysis; 5: moribund or death.

### Microinjections

Microinjection into corpus callosum (CC) was performed on 8-week-old WT mice which were anesthetized and maintained with inhalational isoflurane and oxygen supplemented with buprenorphine given subcutaneously. The cells of interest were injected into the CC with a Hamilton needle at the following coordinate: 1.0 mm rostral, 1.0 mm lateral, and 1.65 mm ventral to bregma, and the needle was left in place for 5 mins following injection to reduce backflow. Microinjection into cerebellar slice proceeded at an angle of 45° with a hand-hold Hamilton needle. The cells of interest were injected into the white matter area of the cerebellar slice, and the slices were cultured for 3 days before immunostaining.

### Drug administrations

<u>Tamoxifen:</u> tamoxifen was dissolved in ethanol/corn oil (1:9) at a dosage of 30 mg/ml. These *Pdgfr*α*^CreER^* mice were given tamoxifen gavage with 3 mg each day for 5 consecutive days.

<u>CD4 depletion antibody:</u> an anti-CD4 monoclonal antibody was given intraperitoneally to EAE mice at a dosage of 400 μg per mouse on 9 dpi, 12 dpi and 15 dpi.

<u>Maraviroc (MVC):</u> to block the effect of CCL4, a CCR5 inhibitor MVC was dissolved in DMSO/corn oil (1:9) and given intraperitoneally to EAE mice at a dosage of 50 mg/kg daily from 7 dpi.

### Primary cell cultures

<u>Mouse primary OPC:</u> OPCs were isolated from postnatal day 8 (P8) mouse cortices by immunopanning as described previously ^12^. Briefly, mouse brains were minced and dissociated with papain and DNase I at 37LJ for 60 mins. After trituration, cells were incubated with primary antibody PDGFRα in a panning buffer at room temperature for 30 mins, then seeded onto pre-overnight-coated secondary antibodies panning dishes for another 30 mins. Purified mouse OPCs were released from the panning dish by 0.05% trypsin and cultured in poly-D-lysine-coated 10 cm dishes or 24-well plates with coverslips for RNA extraction and immunohistochemistry after treatment.

<u>Mouse splenocytes:</u> splenocytes were isolated by disrupting the spleen in RPMI1640 medium and passing through a 70 μm mesh nylon strainer to remove aggregates and debris. The cell suspension was then centrifuged and subjected to Red-blood-cell lysis treatment to obtain purified splenocytes.

<u>Mouse T cells:</u> CD4^+^ T cell, CD8^+^ T cell, and Naïve CD4^+^ T cell were isolated from splenocytes with commercial kits (STEMCELL Technologies, Cat No. 19852, 19853, 19765) according to the manufacturer’s instructions. Th1 cells were induced from isolated Naïve CD4^+^ T cells, the Naïve CD4^+^ T cells were plated on 24-well plates which were pre-coated with 2 μg/ml anti-CD3ε overnight, then culture with RPMI1640 medium containing 10% FBS, 2 μg/ml anti-CD28, 1 μg/ml IL-4 antibody, 5 ng/ml IL-2 and 10 ng/ml IL-12 for 3 days in 5 % CO_2_ incubator at 37 °C.

<u>Mouse peritoneal macrophages:</u> Macrophages were isolated from adult mouse peritoneal cavity as described previously ^81^. Briefly, 3ml 3% Brewer thioglycolate medium was injected into the mouse peritoneal cavity for inflammatory response proceeding. After 3 days, fluid from the mice peritoneum was aspirated and centrifuged to remove the supernatant, and the remaining peritoneal exudate cells were resuspended in RPMI1640 media with 10% FBS and allowed to adhere to substrate by culturing at 37 °C for 2h, after which nonadherent cells were washed and removed to purify adherent macrophage. Macrophages at this stage were processed for analysis or co-cultured with Th1 cells for FACS and immunohistochemistry after treatment.

<u>Conditional media collection:</u> The purified mouse OPCs were plated onto 10 cm dishes and cultured for 3 days. The cultured medium was centrifuged to remove cell debris and then collected as conditional media for downstream experiments or stored at -80°C. The conditional media from *Apc^fl/fl^* OPC cultures was labeled as healthy OPC CM, and the conditional media from *Olig2^Cre^*;*Apc^fl/fl^*OPC cultures was labeled as WNT-OPC CM. Th1 cells and macrophage-derived cells were plated to 24-well plates in RPMI1640 media with 10% FBS, the supernatant was collected 48 hrs later as conditional media after centrifuge.

### Cerebellar Slice Culture

Cerebellar slice cultures were following the protocol as previously described ^12^. Briefly, the cerebellum from the P8 mouse brain was dissected and then cut to 300 μm thick with a McIlwain Tissue Chopper (Campden instruments, Cat No. TC752). The cerebellar slices were transferred to the 0.4 μm Millicell-CM organotypic culture inserts (Millipore, Cat No. PICM0RG50) with slice culture medium containing 50% DMEM, 25% HBSS, 25% heat-inactivated horse serum, 1% N2 and penicillin-streptomycin. Slice were then cultured at 37 °C and 5% CO_2_ for 24 hrs, and injected with cells of interest.

### Immunofluorescence staining

Mice were anesthetized and transcardially perfused with PBS and 4% paraformaldehyde (PFA). The spinal cord and brain were dissected, post-fixed in 4% PFA overnight, and cryoprotected in 30% sucrose. A series of 16-μm sections were obtained from each sample using a cryostat microtome. For coverslip immunofluorescence staining, cells on coverslips were fixed with 4% PFA for 15 mins. Sections or coverslips were blocked with 2% bovine serum albumin (BSA) in 0.5% Triton-PBS for 1 hour at room temperature, followed by primary antibody incubation overnight at 4°C. Primary antibodies were to the following proteins: Rat anti-MBP (Millipore, MAB386), Rabbit anti-SMI32 (Abcam, ab8135), Rabbit anti-RNF43 (Abcam, ab217787), Goat anti-PDGFRα (R&D, AF1062), Rabbit anti-CD4 (Abcam, ab183685), Rat anti-CD4 (Thermo, 14967782), Rabbit anti-CD3 (Abcam, ab5690), Rat anti-CD8 (Thermo, 14008182), Rat anti-CD19 (Thermo, 14019482), Rabbit anti-CCL4 (Abcam, ab45690), Mouse anti-OLIG2 (Millipore, MABN50), Rabbit anti-OLIG2 (Millipore, AB9610), Mouse anti-CC1 (Millipore, OP80), Mouse anti-IP3R2 (Santa Cruz, SC393434), Goat anti-Iba1 (Wako, 011-27991), Goat anti-GFAP (Abcam, ab53554), Rabbit anti-T-bet (Affinity, DF7759), Mouse anti-T-bet (Thermo, 14582580), Rat anti-RORγ(t) (Thermo, 14698880), Mouse anti-Ki67 (BD, 550609), Rabbit anti-Ki67 (Thermo, MA514520), Rabbit anti-dMBP (Millipore, AB5864), Rabbit anti-NG2 (Millipore, MAB5320), Rabbit anti-Cleaved Caspase-3 (CST, 9661), Rabbit anti-Phospho-MLKL (CST, 37333), Rabbit anti-Cleaved C-terminal GSDMD (Abcam, ab255603). Appropriate Alexa Fluor conjugated secondary IgG antibodies were used to visualize the primary antibodies and further counterstained with DAPI. Details of primary and secondary antibodies were listed in the key resources table. EdU staining was performed to detect cell proliferation using the Cell-Light EdU Apollo In Vitro Kit (RIBOBIO) according to the manufacturer’s protocol. Cell apoptosis was detected by TUNEL assay with In Situ Cell Death Detection Kit (Roche) following the manufacturer’s protocol. All the slides were scanned using VS200 Research Slide Scanner (Olympus) or Ixplore SpinSR confocal microscope (Olympus) and processed with OlyVIA software.

### Migration assay

Cell migration was performed using 24 well-format 5.0 μm Transwell inserts (Labselect, Cat No.14331). The experimental conditions were pre-determined in a series of pilot experiments. The cells of interest were added at 50000 cells/well in the upper chambers and were incubated at 37°C for 3 hrs with different medium treatments in the lower chambers. The migratory cells in the lower chambers were enumerated with an automated cell counter. The data are expressed as the mean number of migratory cells in the lower chamber as a percentage of the total number.

### Flow cytometry analyses

Spinal cords, spleens, and blood from the EAE mice were harvested at different disease points as indicated in the main text. Mice were perfused intracardially with cold HBSS under deep anesthesia, Spinal cords were carefully smashed into small pieces and digested with Collagenase (2 mg/ml) and DNase I (1 mg/ml) in RPMI1640 for 30 min in a thermomixer at 250 rpm and 37°C. The cell suspension was passed through a 70 μm cell strainer and mixed with Percoll/HBSS solution, then the mononuclear cell interface was collected over a 30%/70% Percoll gradient centrifuge. The spleen and blood were lysed with RBC lysis buffer to remove red blood cells and collected in PBS with 2% FBS. Single-cell suspensions of the spinal cord, spleen, and blood were stained with fluorescence-conjugated antibodies for 30mins on ice and processed for downstream analysis. For intracellular staining, cell suspensions were treated with a cell stimulation cocktail (Invitrogen, Cat No.00497593) for 4h at 37 °C. After surface marker staining, cells were fixed with 2% PFA for 20min and permeabilized with Perm/Wash buffer (Biolegend, Cat No.421002) for 30mins, incubated with intracellular antibodies for 30mins on ice, then washed and suspended in PBS with 2% FBS for flow cytometry analysis. FACS antibodies used for flow cytometry staining are: BV421 anti-CD3 (Biolegend, 100228), BV605 anti-CD103 (Biolegend, 121433), APC anti-CD103 (Biolegend, 110905), APC anti-CD11b (Biolegend, 101212), BV510 anti-CD11b (Biolegend, 101245), FITC anti-CD11c (Biolegend, 117305), PE anti-CD122 (Biolegend, 105906), BUV737 anti-CD127 (BD, 612841), BV421 anti-CD163 (Biolegend, 155309), BV605 anti-CXCR3 (Biolegend, 126523), PE anti-CXCR3 (Biolegend, 126505), BV650 anti-CD19 (Biolegend, 115541), FITC anti-CD19 (Biolegend, 115506), BV785 anti-CD19 (Biolegend, 115543), PE anti-CCR4 (Biolegend, 131204), APC anti-CCR4 (Biolegend, 131211), PE/Cy7 anti-CCR5 (Biolegend, 107017), BV785 anti-CCR6 (Biolegend, 129823), BV650 anti-CD25 (Biolegend, 102037), PerCP/Cy5.5 anti-CD4 (Biolegend, 100540), BV785 anti-CD4 (Biolegend, 100551), PerCP/Cy5.5 anti-CD44 (Biolegend, 103031), PE/Cy7 anti-CD44 (Biolegend, 103030), PerCP/Cy5.5 anti-B220 (Biolegend, 103236), APC anti-CD45 (Biolegend, 103112), BV786 anti-CD49d (BD, 564397), BV510 anti-CD62L (Biolegend, 104441), BV650 anti-CD86 (Biolegend, 105035), BV785 anti-CD86 (Biolegend, 105043), BV510 anti-CD8a (Biolegend, 100752), BV711 anti-CD8a (Biolegend, 100759), PE anti-F4/80 (Biolegend, 157304), PE/Cy7 anti-Gr-1 (Biolegend, 108416), PE anti-Ly6C (Biolegend, 128007), PE/Cy7 anti-Ly6G (Biolegend, 127617), FITC anti-NK1.1 (Biolegend, 108705), PE anti-NK1.1 (Biolegend, 108708), APC anti-NK1.1 (Biolegend, 108709), FITC anti-IFN-γ (Biolegend, 505806), FITC anti-CD69 (Biolegend, 104505), APC anti-GM-CSF (Biolegend, 505413), PE anti-IL-2 (Biolegend, 503807), BV510 anti-TNF-α (Biolegend, 506339), AF647 anti-FOXP3 (Biolegend, 126407), PE anti-IL-17A (Biolegend, 506903), PE MOG_35-55_ Tetramer (MBL, TS-M7041), eF780 Fixable Viability Dye (Thermo, 65086514). Optimal concentrations for FACS antibodies were determined using titration experiments. Data were collected by BD LSRFortessa flow cytometer and BD FACSAria III flow cytometer. All data were analyzed by FlowJo and plots were gated from singlets, live, and CD45^+^ lymphocytes.

### Automated population identification in FACS data

FACS data for immune cells isolated from 18 dpi EAE mice spinal cord were acquired on BD LSRFortessa flow cytometer, data were compensated and exported into FlowJo for unbiased analysis as described previously ^82^. The data matrix was reduced to the same number and generated a subpopulation containing cells distributed regularly and randomly throughout the selected parent population. Uniform Manifold Approximation and Projection (UMAP) algorithm was used for dimensionality reduction for the visualization of high-dimensional data, the heatmaps display medium expression levels for the markers.

### RNA extraction and RT–qPCR

Total RNA from tissues or cell samples was isolated using the Qiagen RNeasy Mini Kit following the manufacturer’s instructions. Reverse transcription reaction was carried out using the Takara PrimeScript RT Reagent Kit. The qPCR experiments were performed with the FastStart Universal SYBR Mix (Roche, Cat No. 4913914001) and Accurate 96 Real-Time PCR System (DLAB). The ΔΔ*C_T_*method was applied to determine differences in gene expression levels after normalization to the arithmetic mean of GAPDH.

### Bulk RNA sequencing

Bulk RNA sequencing was performed following a previously described EASY-RNA-seq method ^83^. Briefly, OPCs were isolated by immunopanning and RNA was extracted from the isolated cells by Trizol (Thermo, Cat No. 15596026) according to the manufacturer’s protocol. RNA-seq was performed by the Beijing Genomics Institute (BGI). Differential expression analysis was performed on read counts using DESeq2. Pair-wise comparisons were performed between the *Olig2^Cr^*^e^;*Apc^fl/fl^*OPC and the *Apc^fl/fl^* OPC control. Immune cell populations were identified by FACS antibody listed in the key resource table. After gating for singlet live cells, Th1 cell and macrophage-derived cells (CD11b^+^Gr-1^+^NK1.1^+^) were indexed-sorted into Trizol. RNA was isolated by using the Direct-zol™ RNA Miniprep Kit (ZYMO, Cat No. R2050) according to the manufacturer’s instructions. The qualified RNA samples were processed for reverse transcription with SuperScript III reverse transcription kit and second-strand cDNA synthesis with the help of DNA polymerase I and T4 DNA ligase, following the manufacturer’s instructions. Samples were then barcoded following the protocol of TruePrep DNA Library Prep Kit with P5/P7 adapter primers (Vazyme, Cat No. TD503), the amplicon was purified via the VAHTS™ DNA clean beads kit (Vazyme, Cat No. N411-02), and processed for sequencing library preparation. All EASY-RNAseq libraries were sequenced through the Illumina HiSeq platform with a 150-bp pair-end model. Raw reads were passed through adapter filtration and process for potential rRNA contaminants clearance. The clean data were then aligned to GRCm38 for genome reference. Normalization of gene expression was performed by transferring the read counts to FPKM values, and both the read counts and FPKM values were used in visualization.

### Human single-cell RNA-seq data reanalysis

Single-cell RNA-seq data from the previously published dataset was obtained from the Gene Expression Omnibus (GEO) under the accession GSE227781 ^29^. Data integration was performed using Canonical Correlation Analysis (CCA) implemented in the Seurat R package (version 4.2.1) ^84^. Cells with a total count of RNA greater than 50000 and a percentage of mitochondrial genes higher than 20% were filtered out from the downstream analysis to ensure data quality. Principal Component Analysis (PCA) was applied to reduce the dimensionality of the integrated datasets, and the top 20 principal components were retained for further analysis. FindNeighbors and FindClusters functions were implemented to identify the nearest neighbors and obtain cellular clusters, which grouped cells that exhibited similar gene expression profiles, allowing for the identification of biologically relevant cellular populations in the dataset. FindAllMarkers function was used to identify differentially expressed genes (DEGs) for each cellular cluster. The clusters were annotated using canonical cell markers previously characterized and associated with specific cell types. The average expression of genes of interest in OPCs across different conditions was calculated and visualized with heatmaps.

### Quantification and statistical analysis

<u>Quantifications:</u> to quantify positive cell numbers, cell counting was conducted on nine randomly chosen fields for each sample using the Image Pro Plus image analysis system. The areas of interest (AOI) were separated by setting the threshold at least two times the background. Cell counting and fluorescence intensity analyses were conducted on six randomly chosen fields for each sample using the Image Pro Plus image analysis system.

<u>Statistical analysis:</u> statistical significance between groups was determined with GraphPad Prism software. The unpaired *t*-test was used to determine the significance between two experimental groups. One-way analysis of variance (ANOVA) was used to determine the significance among three and four groups. A probability of P<0.05 was considered statistically significant. All significant statistical results are indicated within the figures with the following conventions: \**p* < 0.05, \*\**p* < 0.01, \*\*\**p* < 0.001, \*\*\*\**p* < 0.0001. Error bars represent SEM. No statistical methods were used to pre-determine sample sizes but our sample sizes are similar to those reported in previous publications ^11,85^. Data distribution was assumed to be normal, but this was not formally tested. All experiments were performed at least 3 times, and the findings were replicated in individual mice and cell cultures in each experiment.

### Data and code availability

Bulk RNA-seq data have been deposited at GEO and are publicly available as of the date of publication. Accession numbers are listed in the key resources table. This paper does not report any original code. Any additional information required to reanalyze the data reported in this paper is available from the lead contact upon request.

## Supplemental information titles and legends

**Figure S1.**
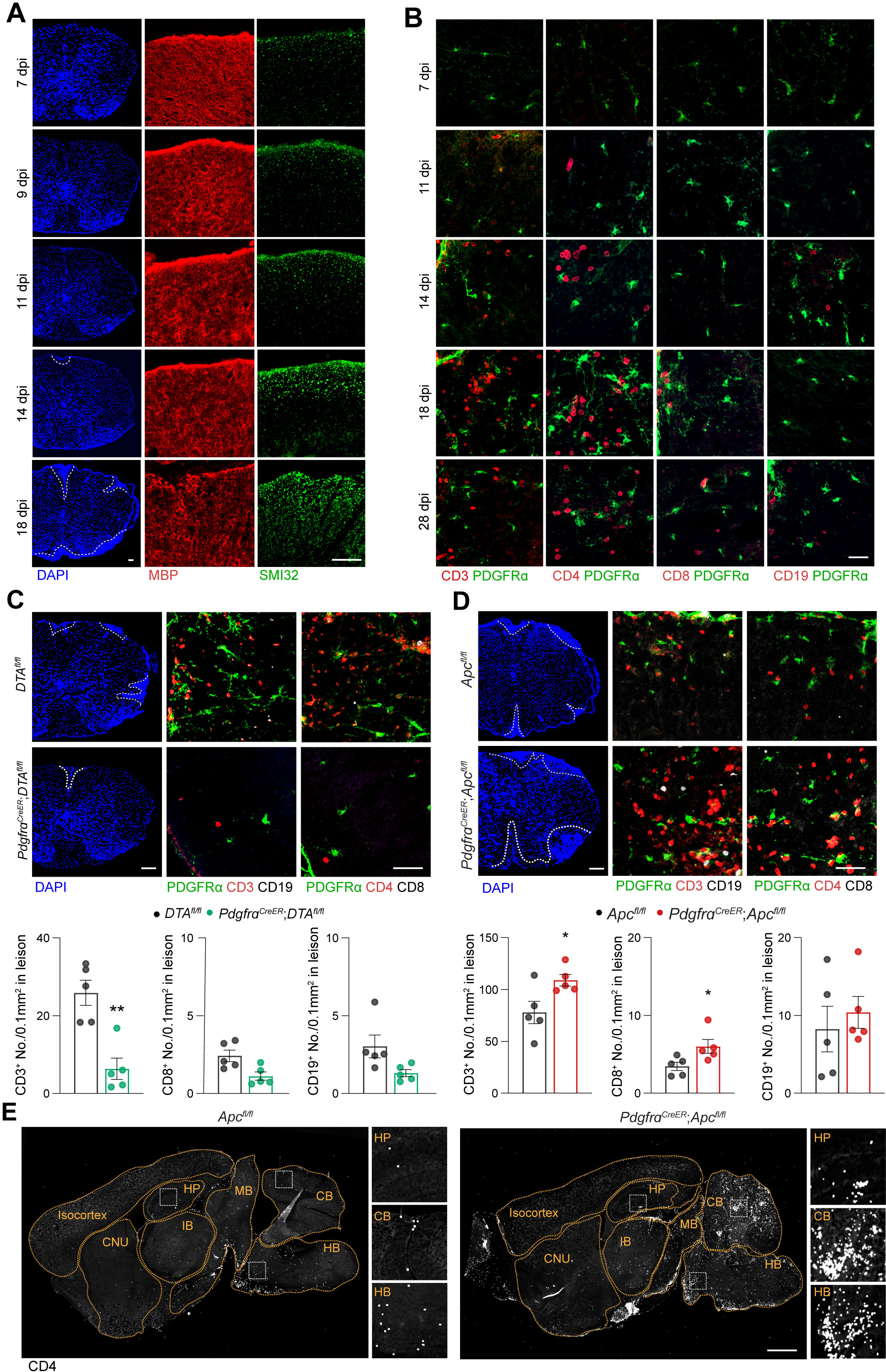
hWnt-OPCs promote EAE pathogenesis and CD4^+^ T cell recruitment in early EAE. (A) Representative immunofluorescent images of the EAE spinal cord show MBP and SMI32 changes during the EAE progression. Scale bar represents 50 μm. (B) Representative immunofluorescent images of the EAE spinal cord show PDGFRα^+^ OPC and immune cells during the EAE progression. Scale bar represents 50 μm. (C, D) Representative immunofluorescent images of the spinal cord show accumulated immune cells at 18 dpi in *DTA^fl/fl^* and *Pdgfr*α*^CreER^*;*DTA^fl/fl^* EAE mice, and *Apc^fl/fl^* and *Pdgfr*α*^CreER^*;*Apc^fl/fl^* EAE mice, and quantifications. Scale bar represents 50 μm. (E) Representative immunofluorescent images of CD4^+^ T cells in the brain regions of *Apc^fl/fl^*and *Pdgfr*α*^CreER^*;*Apc^fl/fl^*EAE mice at 18 dpi. CNU: Cerebral nuclei; HP: Hippocampus; IB: Interbrain; MB: Midbrain; CB: Cerebellum; HB: Hindbrain. Scale bar represents 500 μm. Plots show Mean ± SEM and \**p* < 0.05, \*\**p* < 0.01, n=5 mice/group

**Figure S2.**
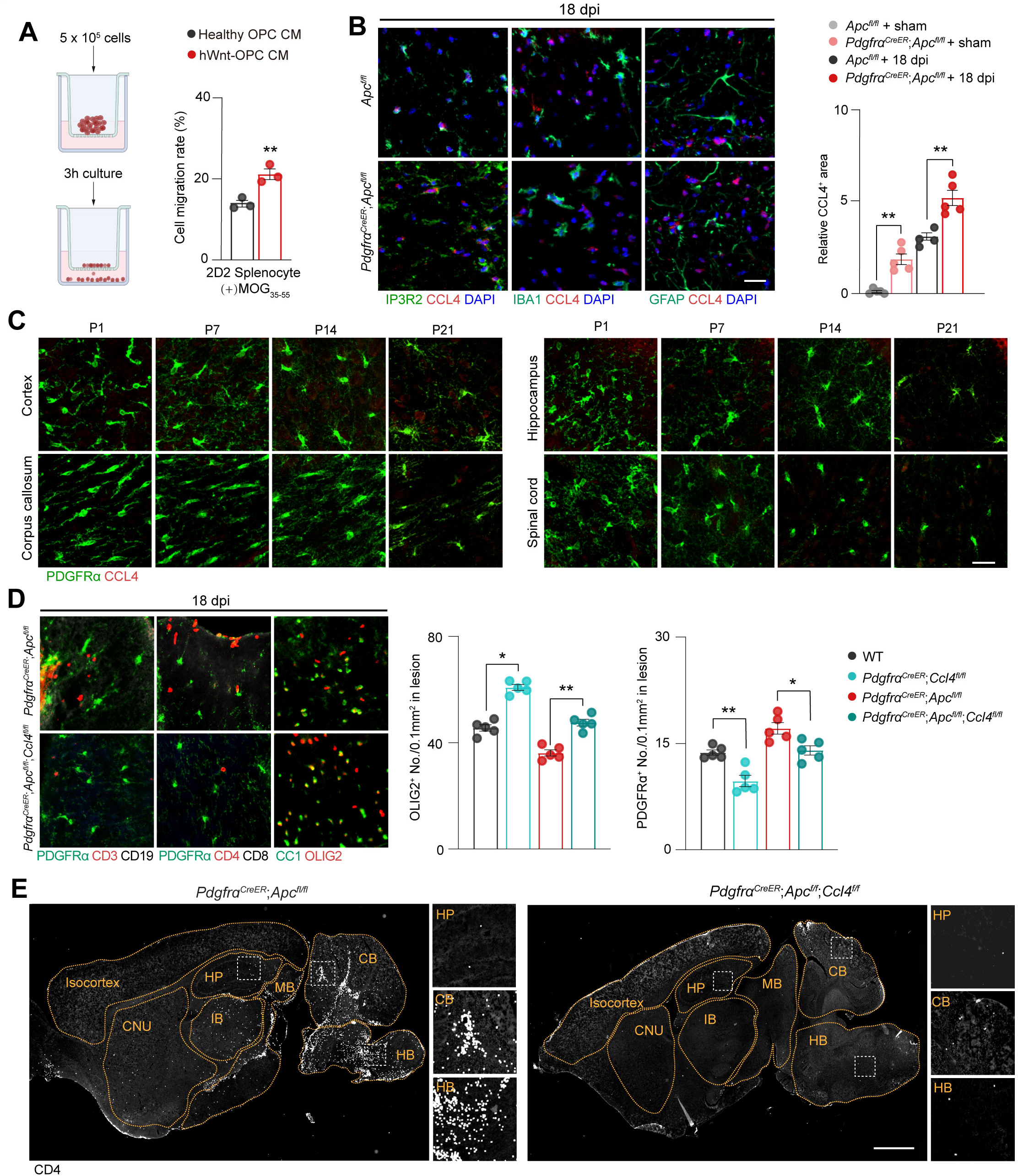
hWnt-OPC-derived CCL4 mediates the CD4^+^ T cell recruitment and EAE pathogenesis. (A) Schematic diagram of Transwell experiment. The cell migration rate of MOG_35-55_ stimulated 2D2 splenocyte in the presence of healthy OPC CM or hWnt-OPC CM. (B) Representative immunofluorescent images of CCL4 and co-staining with glial cell markers and the quantification of *Ccl4* mRNA expression levels in spinal cords of *Apc^fl/fl^* and *Pdgfr*α*^CreER^*;*Apc^fl/fl^* EAE mice at 18 dpi. Scale bar represents 50 μm. (C) Representative immunofluorescent images of CCL4 and co-staining with PDGFRα show no CCL4 is expressed in OPCs at postnatal days. Scale bar represents 50 μm. (D) Representative immunofluorescent images of the spinal cord show accumulated immune cells at 18 dpi and quantifications. Scale bar represents 50 μm. (E) Representative immunofluorescent images and quantification of CD4^+^ T cells in different brain regions of *Pdgfr*α*^CreER^*;*Apc^fl/fl^* EAE mice with or without *Ccl4*-knockout. CNU: Cerebral nuclei; HP: Hippocampus; IB: Interbrain; MB: Midbrain; CB: Cerebellum; HB: Hindbrain. Scale bar represents 500 μm. Plots show Mean ± SEM and \**p* < 0.05, \*\**p* < 0.01, n=5 mice/group otherwise specifically mentioned.

**Figure S3.**
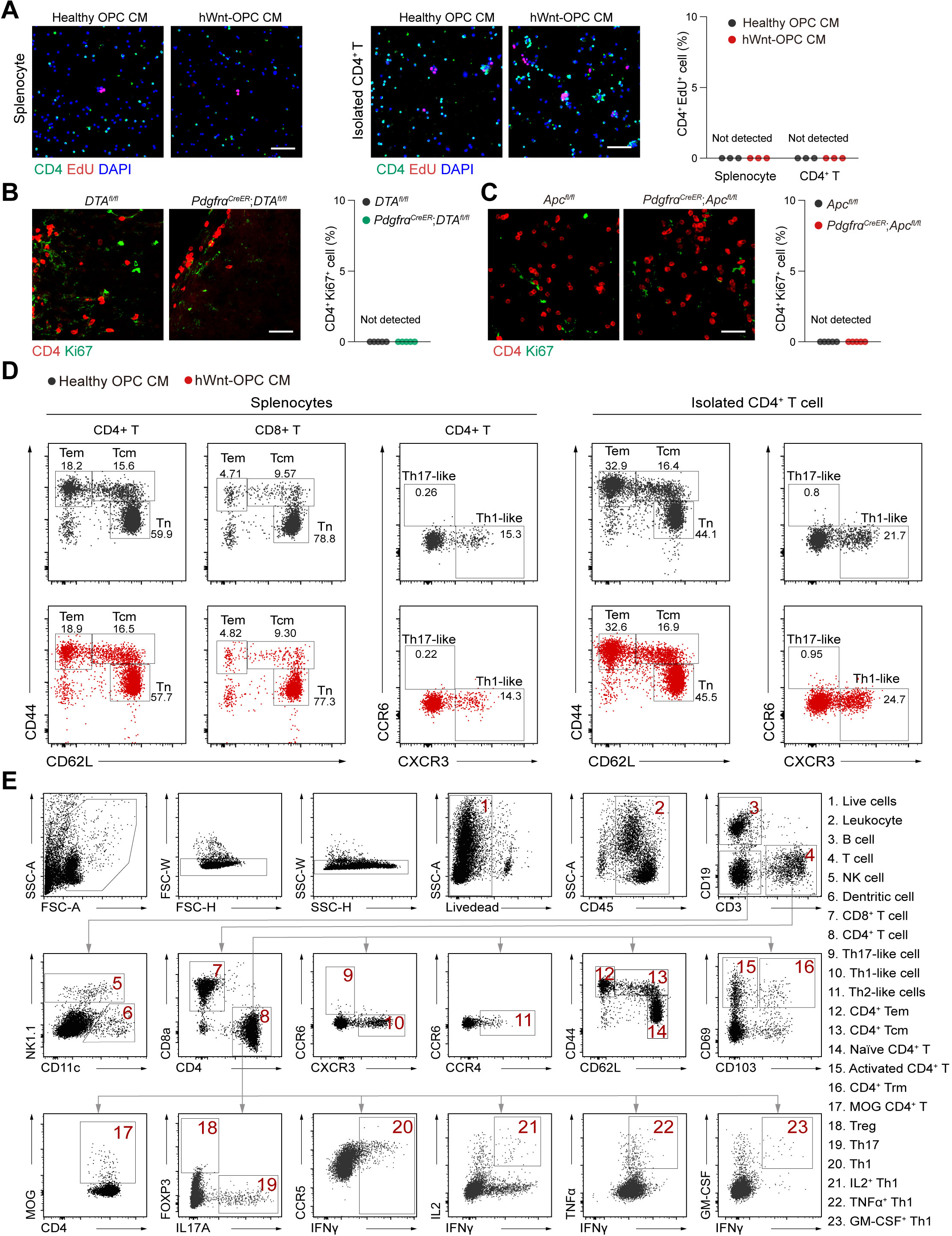
hWnt-OPCs do not promote CD4^+^ T cell proliferation or polarization. (A) Representative immunofluorescent images of CD4 co-staining with proliferation marker EdU and the quantification in splenocyte cultures and isolated CD4^+^ T cell cultures with different OPC-conditional media. Scale bar represents 50 μm. (B, C) Representative immunofluorescent images of CD4 and co-staining with Ki67 show the proliferation of CD4^+^ T cells was not altered in *Pdgfr*α*^CreER^*;*DTA^fl/fl^* EAE mice (B) and *Pdgfr*α*^CreER^*;*Apc^fl/fl^*EAE mice (C) when compared to their littermate control mice. Scale bar represents 50 μm. (D) Flow cytometric analysis of T cells in isolated splenocytes and CD4^+^ T cells when treated with different OPC-conditional media. (E) Flow cytometry gating strategies in this study. Plots show Mean ± SEM, n=3 independent experiments or 5 mice/group otherwise specifically mentioned.

**Figure S4.**
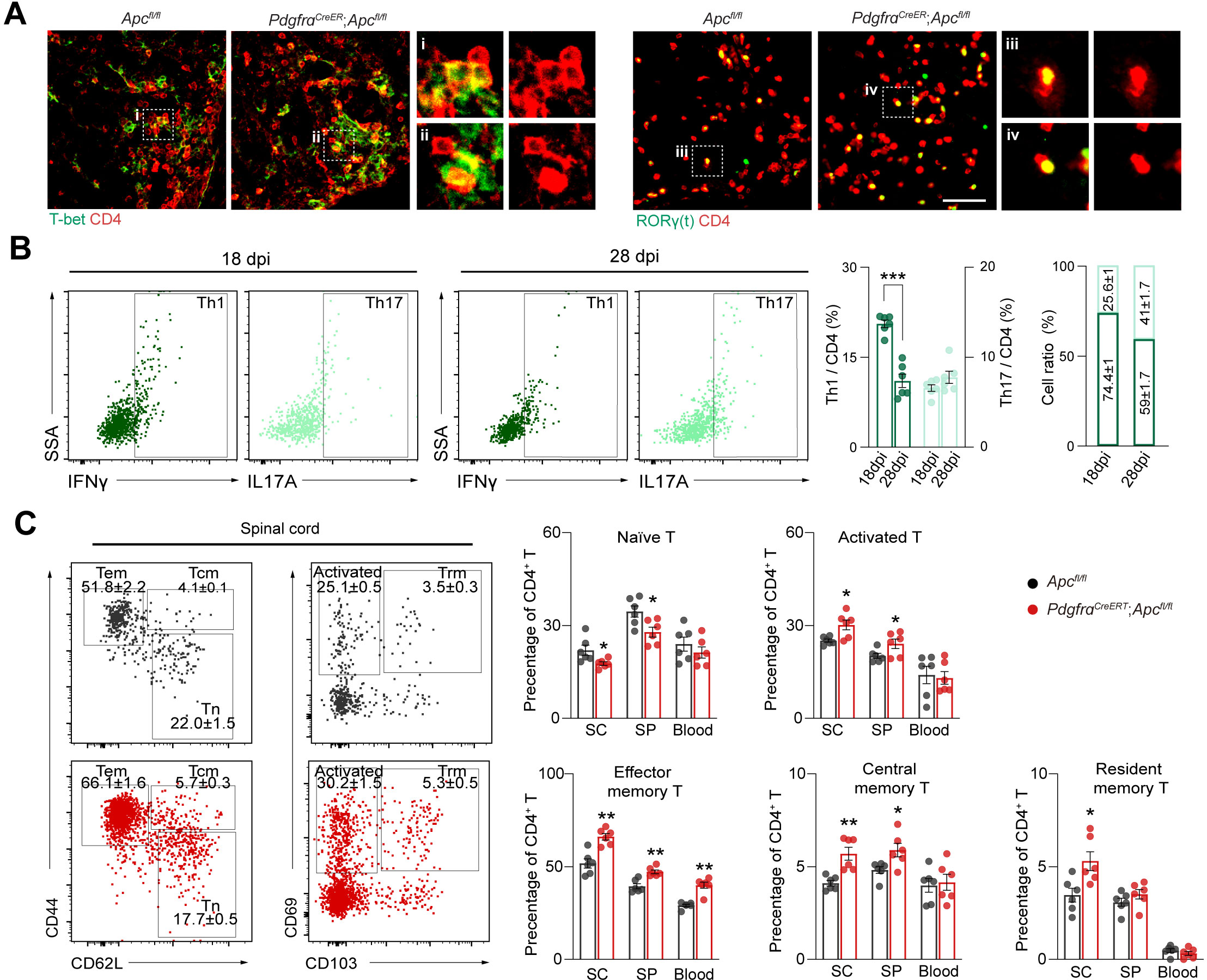
hWnt-OPCs induce Th1 cells in the early EAE. (A) Representative immunofluorescent images of T-bet^+^ and RORγ(t)^+^ CD4^+^ T cells in the spinal cord of EAE mice. Scale bar represents 50 μm (B) Flow cytometric analysis of Th1 and Th17 frequency in EAE mice at 18 dpi and 28 dpi, and the quantification of Th1/Th17 ratio during EAE progression. (C) Flow cytometric analysis of subsets of CD4+ T cells in EAE mice and the quantifications in spinal cord, spleen and blood. Plots show Mean ± SEM and \**p* < 0.05, \*\**p* < 0.01, n=5 mice/group otherwise specifically mentioned.

**Figure S5.**
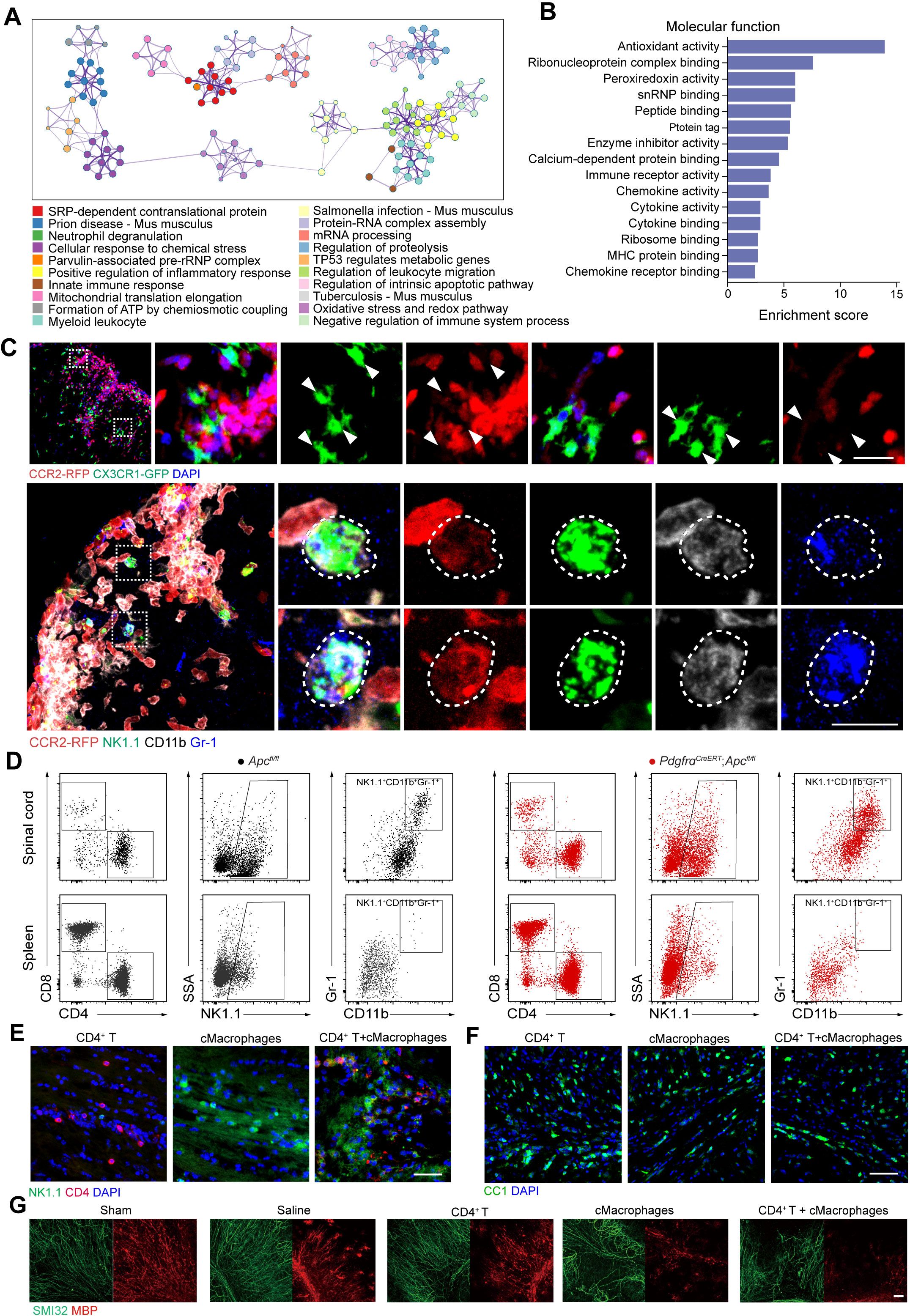
Th1 cells require macrophage-derived cells to cause early demyelination. (A) Network of enriched terms for all highly expressed genes of of NK1.1^+^ CD11b^+^ Gr-1^+^ cells, colored by cluster ID. (B) Bar plot showing the top molecular function pathways associated with the screened highly expressed genes of NK1.1^+^ CD11b^+^ Gr-1^+^ cells. (C) Representative immunofluorescent images in *Olig2^Cre^*;*Apc^fl/fl^*;*Cx3cr1-Gfp*;*Ccr2-Rfp* EAE mice. GFP^+^RFP^+^ co-staining shows the microglia in the CNS and macrophages from the circulation; NK1.1^+^ CD11b^+^ Gr-1^+^ co-staining with RFP shows the co-localization between NK1.1^+^ CD11b^+^ Gr-1^+^ and CCR2-labeled macrophages. Arrowheads point to the cells with only *GFP* expression driven by *Cx3cr1* promoter but not *RFP* signal induced by *Ccr2* promoter. Scale bar represents 50 μm. (D) Flow cytometric analysis of NK1.1^+^CD11b^+^Gr-1^+^ subpopulation in the spinal cord and spleen of *Apc^fl/fl^* and *Pdgfr*α*^CreER^*;*Apc^fl/fl^*EAE mice. (E) Representative immunofluorescent images of CD4 and NK1.1 show the microinjection of CD4^+^ T cells alone, macrophage-derived cells alone, or both cell types combined in the corpus callosum. Scale bar represents 100 μm. (F) Representative immunofluorescent images of CC1 show the demyelination in the corpus callosum after cellular microinjections. Scale bar represents 100 μm. (G) Representative immunofluorescent images of SMI32 and MBP show the demyelination in the cultured cerebellum slices after cellular microinjections. Scale bar represents 100 μm.

**Figure S6.**
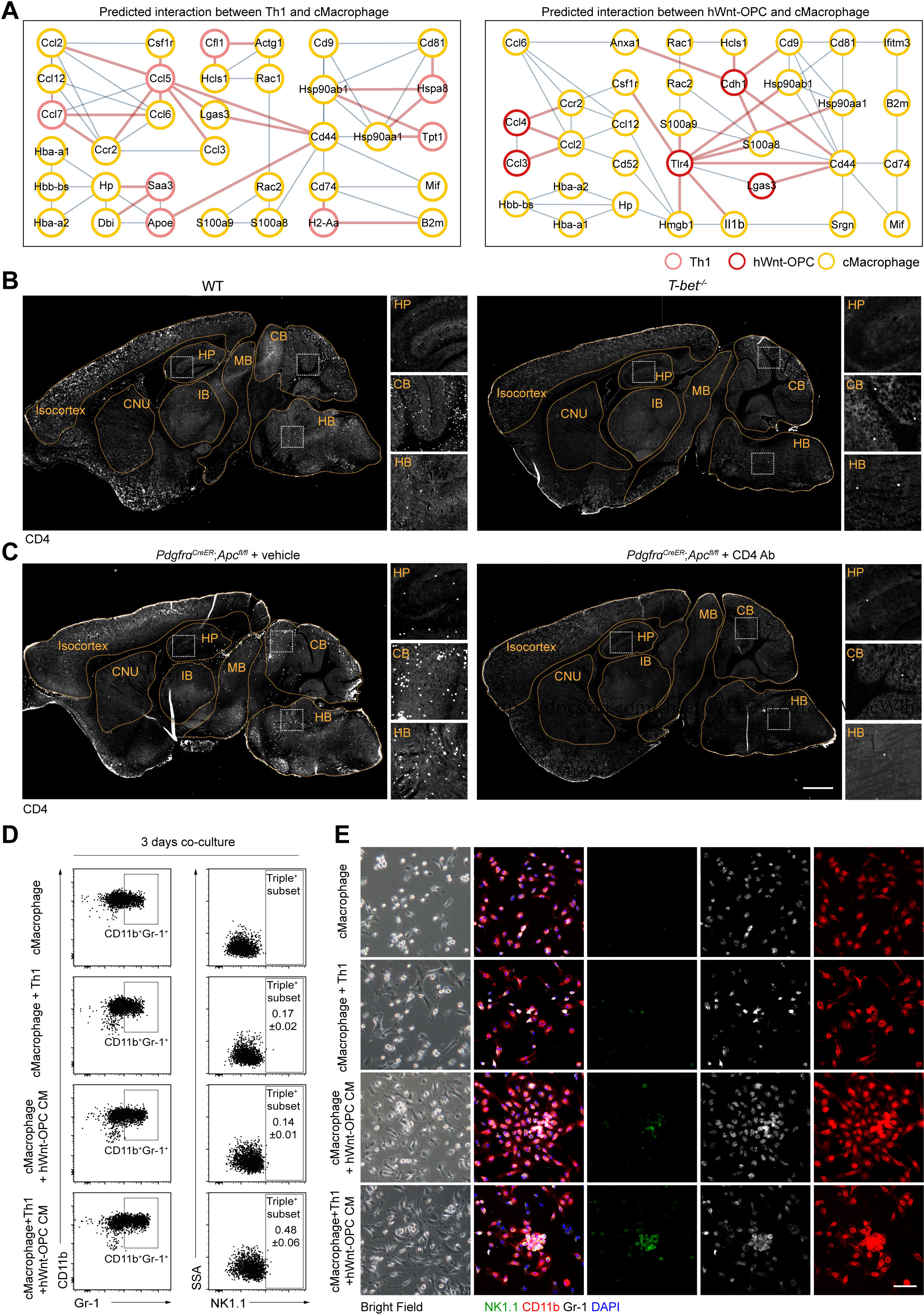
Th1 cells and hWnt-OPCs induce macrophage-derived cells in early EAE. (A) Predicted protein interaction network of Th1, hWnt-OPC and macrophage-derived cells obtained by STRING. (B, C) Representative immunofluorescent images of CD4^+^ T cells in brain regions of *T-bet^-/-^* EAE mice and CD4 depletion antibody-administration EAE mice. CNU: Cerebral nuclei; HP: Hippocampus; IB: Interbrain; MB: Midbrain; CB: Cerebellum; HB: Hindbrain. Scale bar represents 500 μm. (D) Flow cytometric analysis shows 3 days co-culture of primary macrophages and Th1 cells under different conditions. (E) Representative bright field and triple-immunofluorescent staining images for co-culture of primary macrophages and Th1 cells under different conditions.

**Figure S7.**
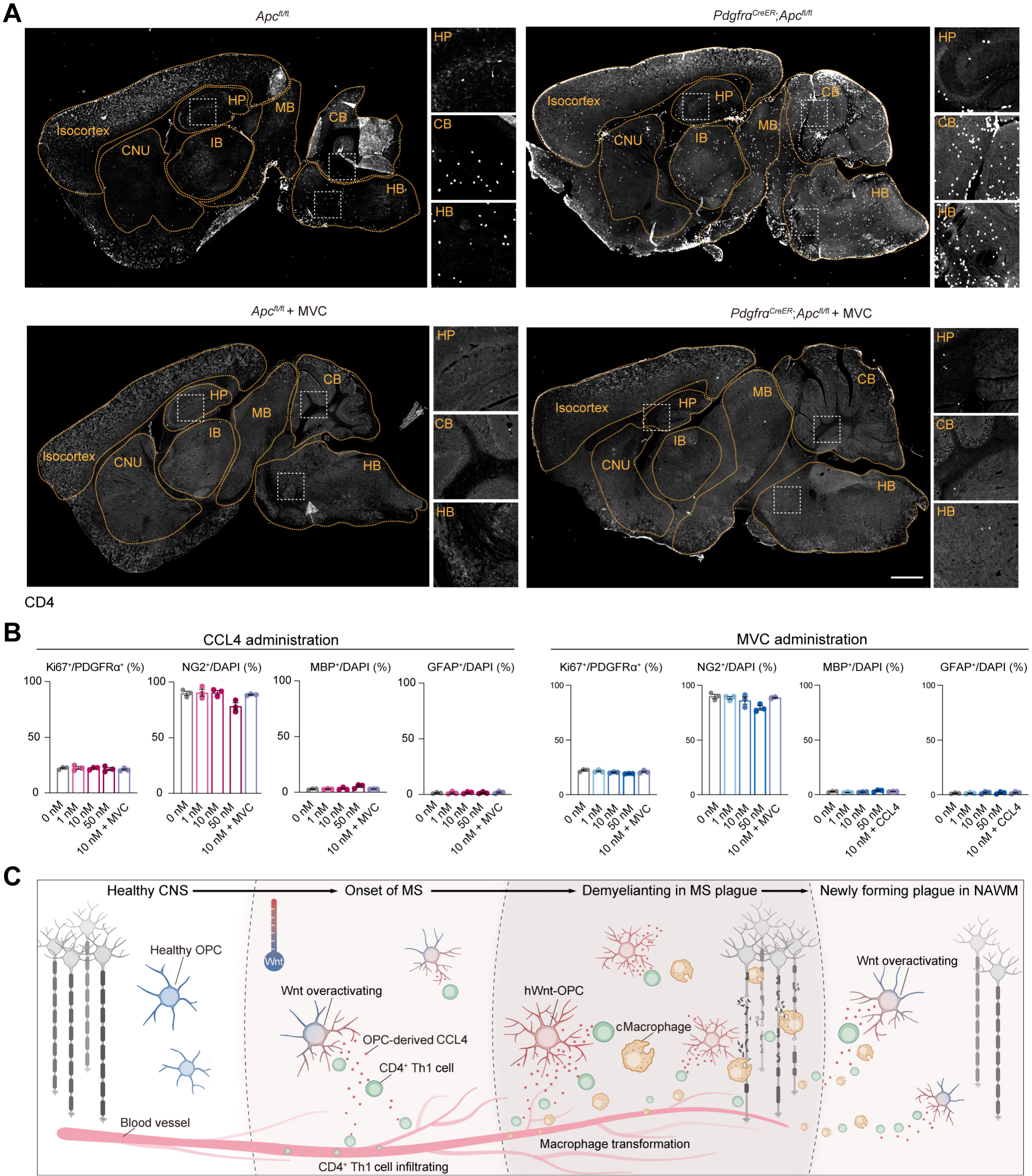
Breaking the OPC-immune cascade by targeting CCL4 attenuates EAE pathologies. (A) Representative immunofluorescent images of CD4^+^ T cells in brain regions of *T-bet^-/-^*EAE mice and CD4 depletion antibody-administration EAE mice. CNU: Cerebral nuclei; HP: Hippocampus; IB: Interbrain; MB: Midbrain; CB: Cerebellum; HB: Hindbrain. Scale bar represents 500 μm. (B) Representative immunofluorescent images and quantifications of proliferation, differentiation of cultured OPCs when treated with or without CCL4 and MVC. Mean ± SEM, n=3 independent experiments. (C) Schematic of WNT-OPC-immune cellular network which may emerge during the onset of the disease or in a newly forming plague in the surrounding normal-appearing white matter (NAWM). In the healthy CNS, healthy OPCs are not affected by immune cells, and myelinated axons are intact. During the onset of MS, OPCs interact with extravasated immune cells, resulting in increased Wnt activation and CCL4 expression in OPCs, and converting them into hWnt-OPCs. Then, hWnt-OPC-derived CCL4 acts on robust Th1 cell infiltration and subsequent cytotoxic macrophage (cMacrophage) infiltration, leading to demyelination and axonal degeneration in the MS plague. With the progression of the disease, infiltrated immune cells interact with healthy OPCs in the NAWM surrounding the MS plague, and a new vicious cycle of hWnt-OPCs and immune cells emerges, recruiting more immune cells in a new location, ultimately leading to progressive tissue damage.

